# Prediction of plant complex traits via integration of multi-omics data

**DOI:** 10.1101/2023.11.14.566971

**Authors:** Peipei Wang, Melissa D. Lehti-Shiu, Serena Lotreck, Kenia Segura Abá, Patrick J. Krysan, Shin-Han Shiu

**Affiliations:** DOE Great Lakes Bioenergy Research Center, Michigan State University, East Lansing, MI 48824, USA; Kunpeng Institute of Modern Agriculture at Foshan, Agricultural Genomics Institute at Shenzhen, Chinese Academy of Agricultural Sciences, Shenzhen, Guangdong 518124, China; Department of Plant Biology, Michigan State University, East Lansing, MI 48824, USA; Department of Computational Mathematics, Science, and Engineering, Michigan State University, East Lansing, MI 48824, USA; Genetics and Genome Sciences Program, Michigan State University, East Lansing, MI 48824, USA; Department of Plant and Agroecosystem Sciences, University of Wisconsin-Madison, Madison, WI 53705, USA

## Abstract

The formation of complex traits is the consequence of genotype and activities at multiple molecular levels. However, connecting genotypes and these activities to complex traits remains challenging. Here, we investigated whether integrating different omics data could improve trait prediction. We built prediction models using genomic, transcriptomic, and methylomic data from the Arabidopsis 1001 Genomes Project for six Arabidopsis traits, and found that transcriptome- and methylome-based models had performances comparable to those of genome-based models. However, when comparing models for flowering time prediction, we found that models built using different omics data identified different benchmark genes. Nine novel genes identified as important for flowering time from our models were experimentally validated as regulating flowering. In addition, we found that gene contributions to flowering time prediction are accession-dependent and that distinct genes contribute to trait prediction in different genetic backgrounds. Models integrating multi-omics data performed best and revealed known and novel gene interactions, extending knowledge about existing regulatory networks underlying flowering time determination. These results demonstrate the feasibility of revealing molecular mechanisms underlying complex traits through multi-omics data integration.

## Introduction

Translating genotypes to phenotype is challenging because the genetic mechanisms underlying trait variation are complex. Although genetic variation information is commonly used to predict phenotypes (i.e., genomic prediction)^1,2^, researchers have had success in using other types of data. For example, transcriptomic data have been used to predict flowering time and yield^3^ and pathogen resistance in plants^4^; methylomic data have been used to predict flowering time and plant height in a panel of epigenetic recombinant inbred lines of *Arabidopsis thaliana*^5,6^; and metabolomic data have been used to predict biomass- and bioenergy-related traits in maize^7^ and yield in rice^8^. Although multi-omics datasets that align with trait variation information are scarce in non-medical, multicellular model systems, the Arabidopsis 1001 Genome Project has generated phenotypic, genomic (G, i.e., biallelic single nucleotide polymorphisms [SNPs]), transcriptomic (T, RNA sequencing), and methylomic (M, gene-body methylation [gbM], or single-site based methylation [ssM]) data for hundreds of accessions of the model plant *A. thaliana*^9,10^. The availability of these datasets provides an opportunity to predict complex traits using machine learning approaches by integrating different data types. Through interpreting these machine learning models, gene features important for prediction of complex traits can be identified to gain a deeper insight into the mechanistic basis of complex traits beyond the few significant quantitative trait loci (QTLs) that can be revealed through genome-wide association studies (GWAS).

Here we generated models for predicting six Arabidopsis traits using G, T, and M data independently and combined (a flow chart illustrating the steps in this study is shown in **Fig. 1**). The six traits, namely flowering time (days until the first flower was open), rosette leaf number (RLN), cauline leaf number (CLN), diameter of the rosette (DoR), rosette branch number (RBN), and stem length (SL), were collected for 383 Arabidopsis accessions from published studies^9–11^ (**Fig. 1a**, **Table S1**). Samples for G, T, and M were taken from mixed rosette leaves harvested just before bolting at 22 °C, flowering time was measured at 10 ℃, and the other five traits were measured at 16 ℃. To obtain a rough estimate of how well the trait variation can be reflected by omics data variation, we first compared the omics similarity matrices among Arabidopsis accessions with the trait similarity matrices (**Fig. 1b**). To establish the predictive models, two algorithms, ridge regression Best Linear Unbiased Prediction (rrBLUP)^12^ and Random Forest (RF)^13^, were used (**Fig. 1c**). In a previous study we found that, despite their simplicity, rrBLUP and RF outperformed other commonly used algorithms for most species and traits tested^14^; furthermore, RF has the advantage of allowing interpretation of the resulting models, in particular allowing identification of non-linear interactions between predictors. To better interpret the predictive models, we compared the genes important for trait prediction with 426 benchmark flowering time genes downloaded from FLOR-ID (http://www.phytosystems.ulg.ac.be/florid/)^15^ and TAIR (https://www.arabidopsis.org/) (**Fig. 1d, Table S2**). Finally, feature interactions were investigated by interpreting the integrated models built using all the G, T, and M features for the benchmark genes (**Fig. 1e**).

**Fig. 1.**
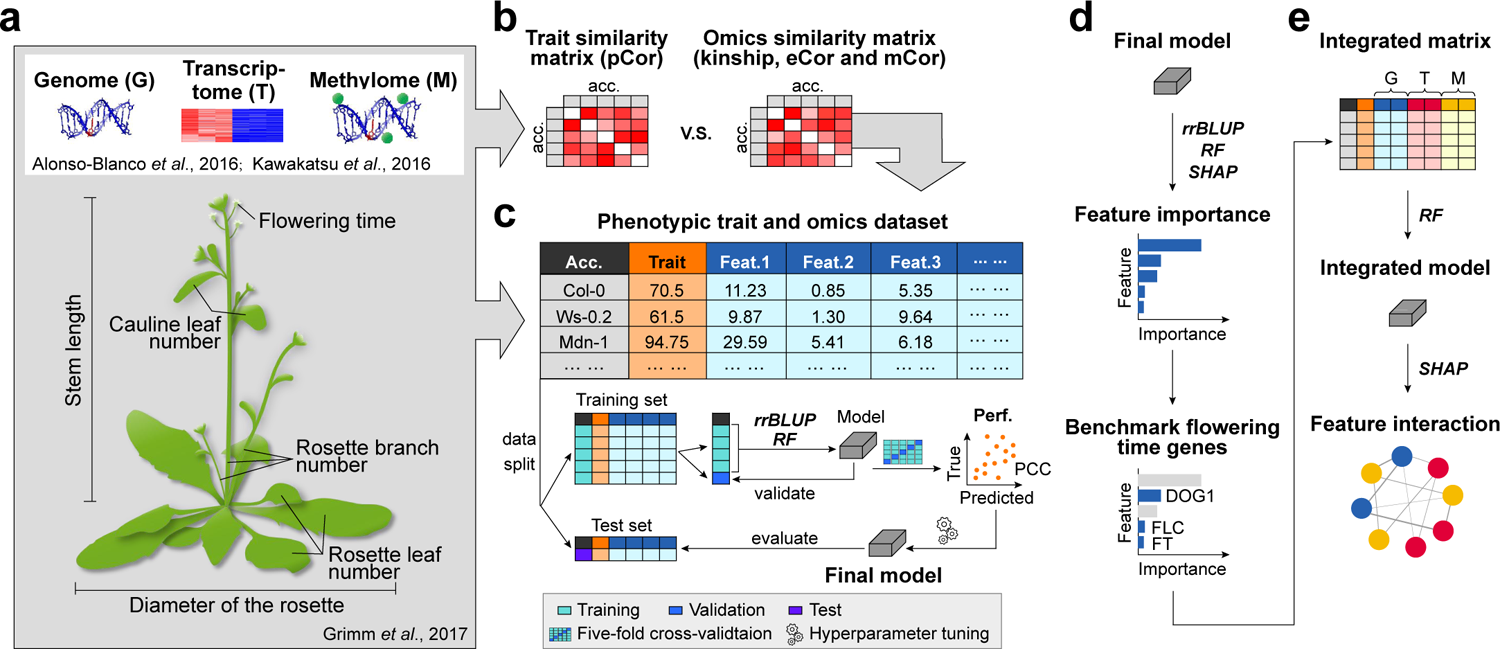
Flow chart of our methodology. (**a**) Three types of omics data (genome [G], transcriptome [T], and methylome [M]) and phenotypic data for six traits were used in this study. Similarities of trait values and omics data between accessions were calculated to produce the trait similarity matrices (pCor) and omics similarity matrices, respectively (**b**). Kinship, eCor (transcriptomic [expression] similarity) and mCor (methylomic similarity) were derived from G, T and gbM (gene-body methylation) data, respectively, and were compared with pCor. (**c**) The original omics data and omics similarity matrices were used as features to build machine learning models for the prediction of six traits with two algorithms: ridge regression Best Linear Unbiased Prediction (rrBLUP) and Random Forest (RF). Each dataset was split into training (80%) and test (20%) sets, and the training set was used to train the models via a five-fold cross-validation scheme and hyperparameter tuning. The final model with optimal hyperparameters was applied to the test data, and the correlation (Pearson Correlation Coefficient [PCC]) between true and predicted trait values for accessions in the test set was measured to evaluate the model performance. (**d**) The final model was further interpreted using the rrBLUP, RF, and SHapley Additive exPlanations (SHAP) approaches to obtain the feature importance values. The features important for flowering time, rosette leaf number, and cauline leaf number were compared with benchmark flowering time genes. (**e**) G, T, and M features of benchmark genes were integrated to build a new model using RF, and the new model was interpreted using SHAP to obtain the interactions between features. Acc.: accessions; Feat.: feature; Perf.: performance.

## Results & Discussion

### Prediction of complex traits using individual multi-omics data

Before investigating the utility of different omics data for plant complex trait prediction, we first examined the omics data structure among accessions (**Fig. 1b**), with the assumption that accessions with more similar trait values are expected to have more similar genetic information (i.e., G, T, or gbM). However, G, T, or gbM data alone explained only a small amount of variation, as there was no obvious relationship between the trait and omics similarity matrices: the correlation (Pearson Correlation Coefficient [PCC]) between phenotypic trait similarity among accessions (pCor; e.g., pCor_flowering time_ see **Fig. S1a**) and the corresponding similarity of G (kinship, **Fig. S1b**), T (eCor, **Fig. S1c**), and gbM (mCor, **Fig. S1d**) only ranged from −0.01 to 0.17 (**Fig. 2a**). This weak correlation is consistent with the findings in our previous study of yield, height, and flowering time in maize^3^, and is expected because only a subset of G/T/gbM variants (e.g., a few SNPs) are expected to contribute to the variation in a complex trait, and the linear correlation between the whole set of variants (e.g., all SNPs) and complex trait values is low. In contrast, kinship and mCor were correlated with each other at a higher level (PCC=0.43, **Fig. 2a**), indicating that gene-body methylation is more heritable than phenotypic traits, or that the M data were confounded by G, which will be discussed later on.

**Fig. 2.**
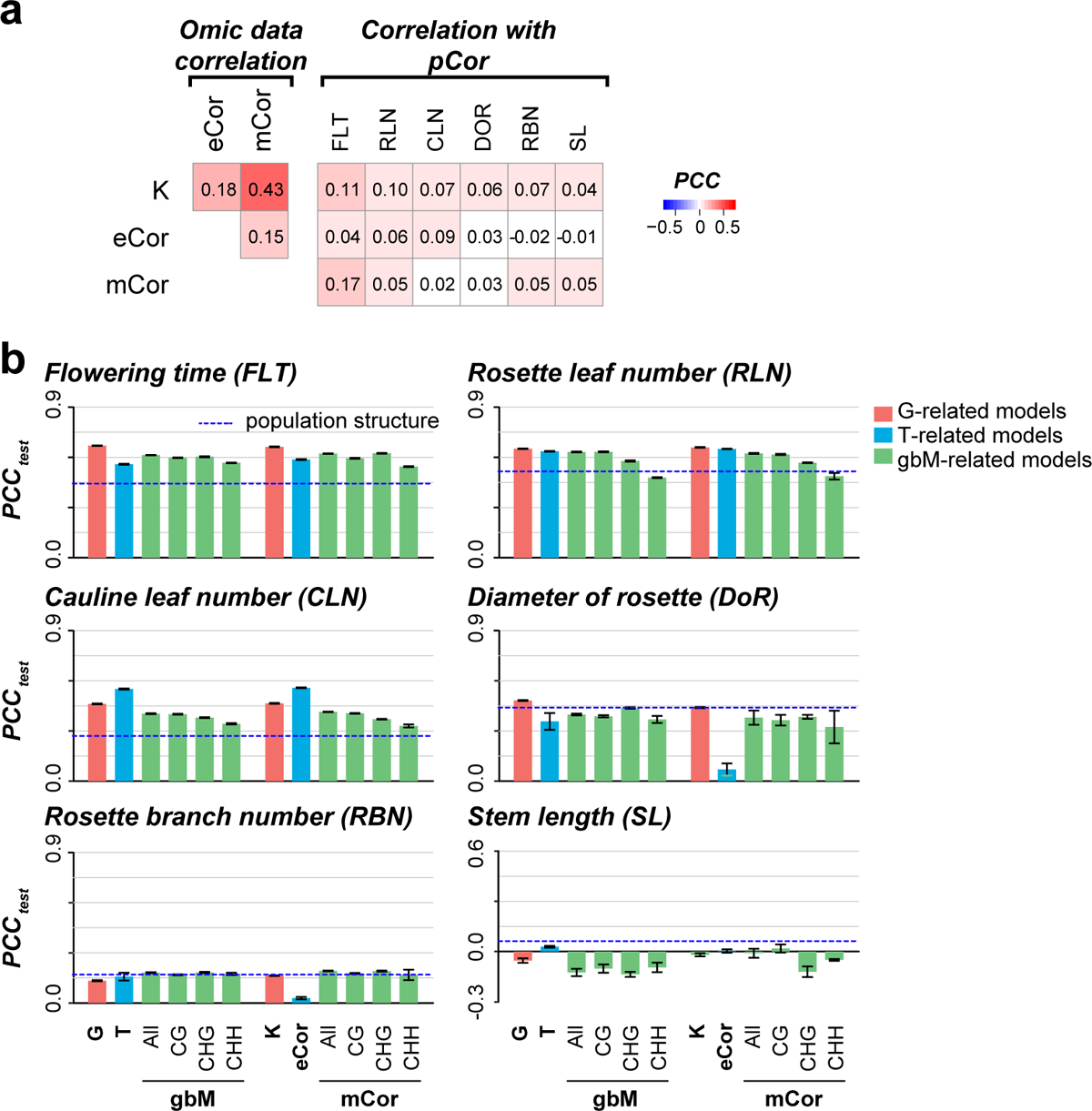
Relationships between omics data and traits, and trait prediction model performance. **(a)** Correlations between kinship, eCor, mCor, and pCor matrices. PCC: Pearson correlation coefficient between omics similarity matrices. (**b**) Performance of prediction models for six traits. Models were built using genomic (G), transcriptomic (T), gene body methylation (gbM) data, or omics data similarities matrices (K, eCor, mCor) with the ridge regression Best Linear Unbiased Prediction (rrBLUP) algorithm (for models built with Random Forest, see **Fig. S2a**). The PCC between true and predicted trait values on the test set (20%, held out before training the model) was used to evaluate model performance. The PCCs of models built using population structure (first five principal components of genetic variation) are indicated by blue dashed lines. Error bars: standard deviation of 10 prediction runs. Colors: models built using G-(red), T-(blue), and gbM-related (green) features. K: kinship; eCor: expression similarity; mCor: methylomic similarity; pCor: phenotypic similarity; G: genomic data; T: transcriptomic data, gbM: gene body methylation data; All: all three types of gene body methylation data, namely, CG-, CHG-, and CHH-types.

While the overall correlations are low, the likelihood that predictive information is encoded within the overall G/T/gbM data led us to build machine learning models to take advantage of all features from single omics data for trait prediction. We established single omics data-based trait prediction models using rrBLUP^12^ and RF^13^, and the model performance was assessed using a hold-out test dataset and measured as the PCC between true and predicted trait values (see **Methods and Fig. 1c**). In addition to the whole set of features for each omics data, the similarity matrices of omics data (i.e., kinship, eCor and mCor, which are derived from G, T, and gbM, respectively) were also used to build models. As expected, the higher the correlation between the omics data and trait values (**Fig. 2a**), the higher the model performance was for most traits and omics data types (**Fig. 2b**, **Fig. S2a**-**d**). For each trait, model performance was similar regardless of algorithm or whether omics data values or value similarities among accessions were used as predictive features (**Fig. 2b, Fig. S2a,e**). Most importantly, G-, T-, and gbM-based models had similar performances (**Fig. 2b, Fig. S2a**), consistent with the findings of our previous maize study using both G and T data for trait prediction^3^.

Since CG, CHG, and CHH methylation have different regulatory mechanisms and functions in plants^16^, we also established models using the different gbM types as separate features. Models using individual gbM data types tended to have similar or poorer performances than models using combined gbM data, and models using CHH methylation tended to have the worst performance (**Fig. 2b, Fig. S2a**). Because of the complexity of M data (e.g., heterogeneity in numbers of methylated cytosine sites within genic regions), we also explored six additional derived M features (single site-based M, hereafter referred to as ssM, see **Methods**) for predicting flowering time as an example. rrBLUP models using ssM features had higher prediction accuracies than those using gbM (**Fig. S3a,b**), but the prediction accuracies for RF models were not higher for unknown reasons (**Fig. S3c,d**). The improved prediction of ssM-based rrBLUP models may be because the ssM features captured the distribution of methylation across each gene, which provided more detailed information about methylation patterns than the gbM data. Taken together, these results indicate that, similar to G and T data^3,17^, M data are also useful for predicting plant traits, and that M data need to be represented in different ways to maximize their predictive power.

### Distinct contributions of omics data to complex trait predictions

To identify the informative variants embedded in the G/T/gbM data, we investigated the importance of features for trait prediction by interpreting the prediction models. Three measures were used to evaluate feature importance: (1) coefficients of features in the rrBLUP models (**Fig. S4a-c**); (2) gini importances in the RF models (**Fig. 3a-c**); and (3) the average absolute SHapley Additive exPlanations (SHAP) values^18^ obtained from RF models (**Fig. S4d-f**). Here, we focus on features important for predicting flowering time (for feature importances for the other five traits, see **Table S3-S8**) because there is abundant knowledge about the genetic control of this trait, which is crucial for interpreting the important features. To allow comparison with T and gbM features, which are gene-based, G variants were mapped to genic regions (see **Methods**). Genes corresponding to or harboring important G/T/gbM features (defined as those with >95^th^ percentile importance values) were considered important for flowering time prediction and are hereafter referred to as important genes. We found weak or no correlation between importance scores from models built using different types of omics data, regardless of the feature importance measure examined (ρ: −0.07– 0.11), and there was little overlap of important genes between models (**Fig. 3, Fig. S4a-f**).

**Fig. 3.**
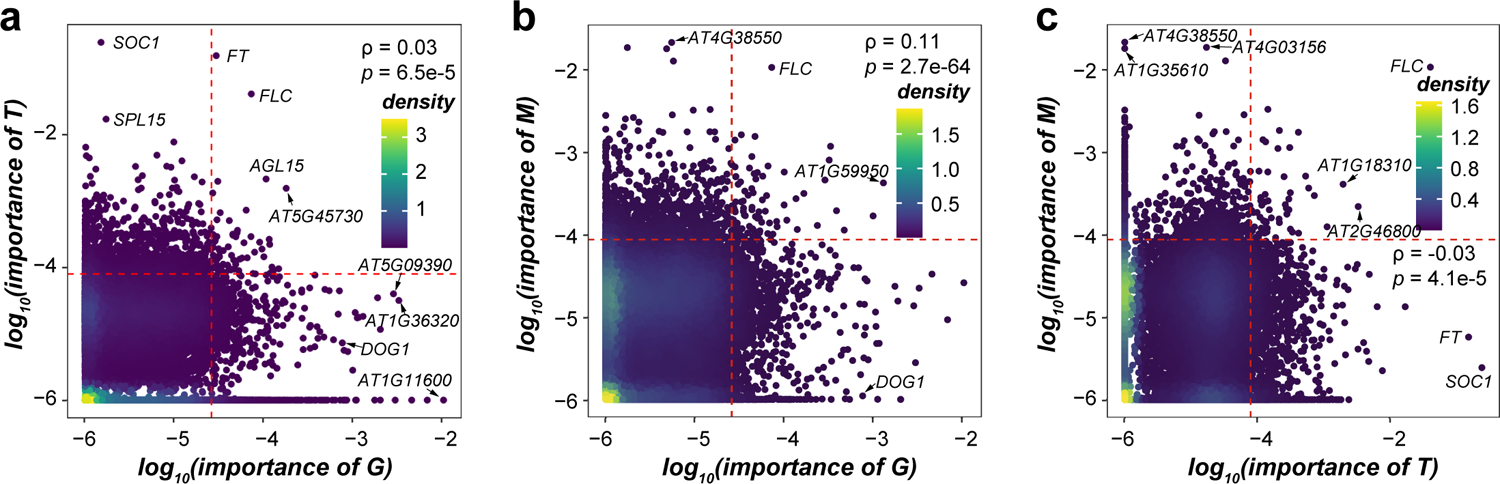
Correlation of feature importance between different types of omics data. **(a-c)** Density scatter plots showing the correlation of the importance scores between genomic (G), transcriptomic (T), and methylomic (M) features of genes when RandomForest gini importance scores were used as measures of importance. (**a**) G vs T. (**b**) G vs M. (**c**) T vs M. All feature importance values < 10^-6^ were assigned a value of 10^-6^. Red dashed line: 95^th^ percentile of feature importance. Density plot color: gene density. ρ: Spearman’s rank correlation coefficient.

These results suggest that the similar trait prediction accuracy of models built with different omics data is not due to shared features, consistent with the findings of the maize study^3^. However, the correlation between the importance of G and gbM variants (**Fig. 3b, Fig. S4b,e**) was higher than that for other comparisons, consistent with the relatively higher correlation between G and gbM similarities across accessions (i.e., kinship and mCor, PCC=0.43, **Fig. 2a**) and suggesting potential confounding effects of G on gbM data. To disentangle gbM from G data, we built trait prediction models with the mCor residuals to exclude the kinship effects (**Methods**). Consistent with a confounding effect of G on gbM, these new models had significantly lower performance compared with models based on the mCor matrix for all traits (differences in PCC scores, which were used as a measure of prediction accuracy: 0.06–0.18, *p*<0.001) except RLN and CLN (differences: 0.03 and 0.01, *p=*0.05 and 0.09, respectively, **Fig. S4g**). However, mCor-based models with confounding effects of kinship removed still performed better at predicting flowering time and RLN than those using the first five principal components to approximate population structure (**Fig. S4g**, the performance of the population structure-based model is used as the baseline for trait prediction). Thus, the components of gbM independent from G are also important for trait prediction.

### Benchmark flowering time genes identified as important features in flowering time prediction models

In the previous section we showed that models built using different omics data identified different important genes (**Fig. 3, Fig. S4a-f**). We next asked how many genes with known functions in flowering time were identified as important genes. Out of 426 benchmark flowering time genes (**Table S2**), 169 were identified as important according to at least one of the three importance measures for at least one of the three individual omics datasets (**Fig. 4**, **Table S9**). Only two genes, *FLOWERING LOCUS C* (*FLC*) and its paralog *MADS AFFECTING FLOWERING 2* (*MAF2*), were identified as important by all three independent omics datasets (blue, **Fig. 4**). This is consistent with the roles of *FLC*^10,19,20^ and *MAF2*^21,22^ in flowering time regulation being established through studies of genetic variation, transcript levels, and methylation levels. Another 27 genes (green, **Fig. 4**) were identified by two independent omics datasets. For instance, *FLOWERING CONTROL LOCUS A* (*FCA*), which increases H3K4 dimethylation in the central region of *FLC* and regulates its expression^23^, was considered important in the G and gbM models. The remaining 140 genes were dataset specific (**Fig. 4**, **Table S9**), such as *SUPPRESSOR OF OVEREXPRESSION OF CO 1* (*SOC1*), which was only identified as important in T models. This is consistent with our observations that there is little overlap between important genes identified by G, T, and gbM models (**Fig. 3, Fig. S4a-f**), and that gbM data, although it is confounded with G information (**Fig. S4g**), can make unique contributions.

**Fig. 4.**
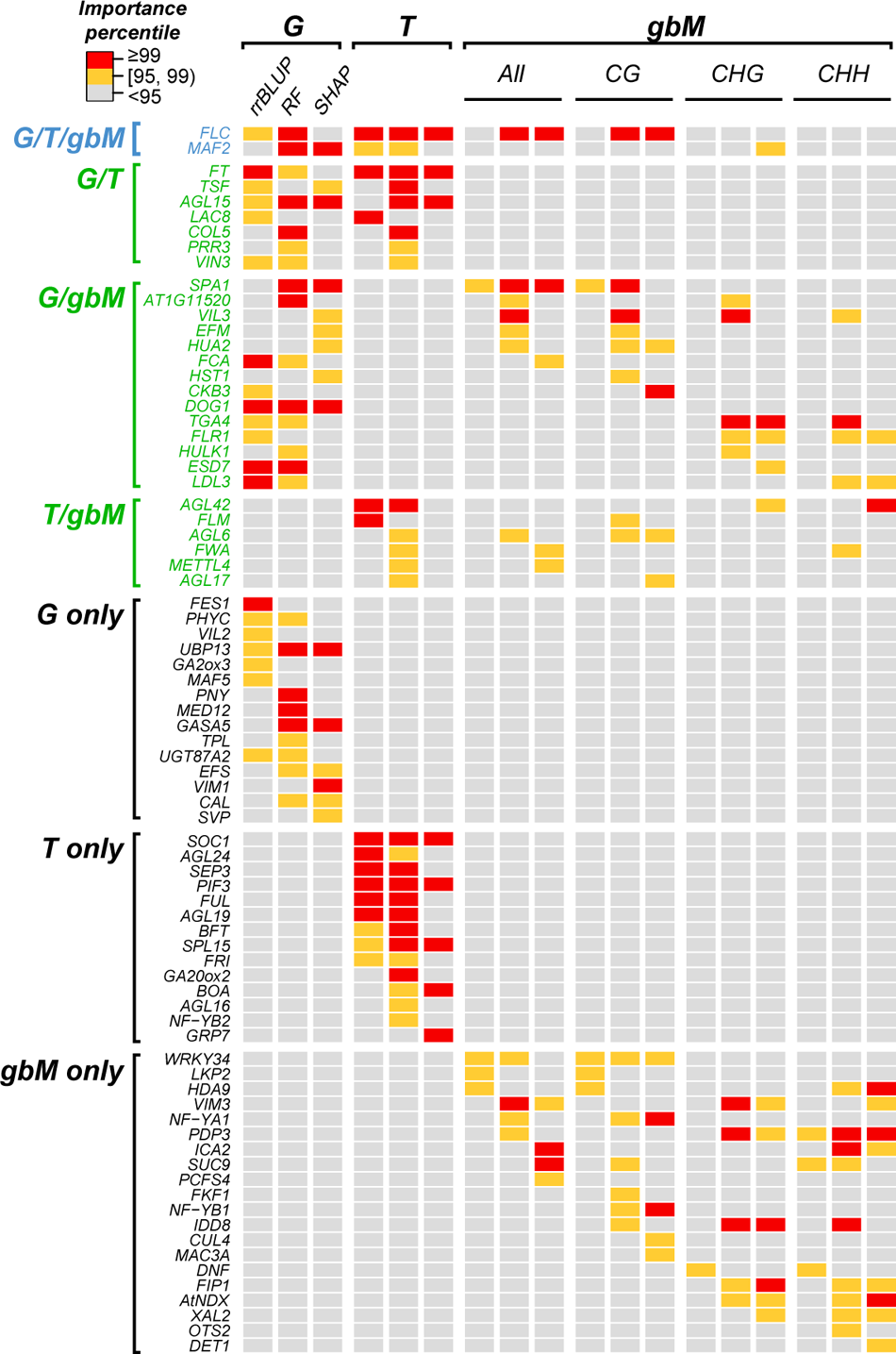
Example benchmark flowering time genes identified using different importance measures and datasets. The heatmap shows how often benchmark flowering time genes were identified as important using different datasets. For each omics dataset, there are three feature importance measures: (1) RF - Random Forest gini values; (2) rrBLUP - absolute value of rrBLUP coefficient; and (3) SHAP - SHapley Additive exPlanations values. Color in the heatmap indicates importance value percentile: ≥99^th^ percentile, red; ≥95^th^ and <99^th^ percentile, orange; <95^th^ percentile, gray. Font color indicates the number of omics datasets that identified a benchmark flowering time gene as important: all datasets, blue; two datasets, green; a single dataset, black. G, T, and M: genomic, transcriptomic, and methylomic data, respectively.

We next asked whether significantly more benchmark flowering time genes than random chance were identified by our models. When the 95^th^ percentile was used as the threshold, only G and T models with RF gini importance scores identified significantly higher proportions of benchmark genes (8.84% and 8.06%, respectively) than expected by random chance (*p=*0.002 and 0.014, respectively, Fisher’s exact test. **Table S10**). To understand why no more benchmark genes were identified than expected by random chance for most models and importance measures, we explored the following potential factors (for the rationale and analysis process, see **Methods**): cut-off threshold (95^th^ or 99^th^ percentile), the number of accessions used for training models, and difference in gene contributions to flowering in short days (SDs) and/or long days (LDs). The former two factors had little impact on the identification of benchmark genes (**Table S10-S12**), except for SHAP values when the 99^th^ percentile was used, where significantly more benchmark genes than expected by random chance were identified for the G and T models (**Table S11**). In addition, genes that when mutated had flowering time phenotypes in two conditions (SDs and LDs) were more likely to be identified using our approaches than those that only had a loss-of-function phenotype in a single condition (SDs or LDs) (**Table S10-11**).

One thing worth noting is that gbM-based models identified no more benchmark genes as important than expected by random chance, regardless of the threshold used (**Table S10-S12**). This may be because gbM does not adequately represent the methylation profiles of a given gene. To assess this, we also interpreted ssM-based models in which the methylation profile across a gene region was represented (see **Methods**). An additional 144 benchmark flowering time genes were recovered as important by at least one ssM feature type/feature importance measure combination (**Table S9**), and more benchmark genes were identified as important for a single ssM model (a maximum of 44 genes, **Table S13**) than a single gbM model (a maximum of 24 genes, **Table S11**). One example gene is *CLEAVAGE STIMULATION FACTOR 77* (*CSTF77*), which is involved in the 3’ processing of antisense *FLC* mRNA^24^. Accessions with CG-type methylation at two different sites in *CSTF77* had significantly longer flowering times than those without (**Fig. S5a,b**), but there were no significant differences in flowering time between accessions with different SNP alleles (**Fig. S5c,d**). In addition, there was no correlation between the expression levels or gbM levels of *CSTF77* and flowering time (**Fig. S5e-h**). These results explain why *CSTF77* was uniquely identified by ssM-based models (**Table S9**).

Another potential reason for the low degree of enrichment of identified benchmark genes for most models is the fact that the plants scored for flowering time phenotypes were grown at 10 °C and those used for the G, T, and M data were grown at 22 °C. We also built models using the same G/T/gbM data to predict flowering time phenotypes recorded at 16 °C^10^, and found that the temperature at which flowering time was scored affected gene importance for flowering time prediction (**Table S14,S15, Fig. S6a**). For example, the expression of *FRIGIDA* (*FRI*), which, when functional, confers a vernalization requirement^25^, had a SHAP value of zero for flowering time prediction at 10 °C but a SHAP value of 0.075 (ranked 14^th^) at 16 °C. This is consistent with the larger differences in flowering time between accessions with functional and non-functional *FRI* copies at 16 °C than at 10 °C (a temperature at which the vernalization requirement is met for accessions with functional *FRI*); the larger SHAP values for *FRI* in the 16 °C model indicate that it makes a higher contribution to flowering time prediction when the plants are not vernalized (**Fig. S6b**). Furthermore, the higher importance rank of *FRI* features at 16 °C compared with 10 °C was also observed for all G- and T-models regardless of the importance measure examined (**Table S14,S15, Fig. S6c,d**).

Since a number of benchmark flowering time genes were identified when RLN and CLN were used as proxies for flowering time, we also asked how many benchmark flowering time genes could be identified by the RLN and CLN models. When only G-, T-, and gbM-based models were examined, 42 genes were identified as important by all flowering time, RLN, and CLN models (**Table S16**); these are generally hub genes in the flowering time regulation network, such as *FT*, *FLC*, *SOC1*, *FRI*, *BROTHER OF FT AND TFL1* (*BFT*), and *SHORT VEGETATIVE PHASE* (*SVP*)^26^.

Consistent with the importance of *FRI* for predicting flowering at 16 °C, *FRI* was important for predicting RLN and CLN in both G- and T-models, probably due to the same temperature (16 °C) at which RLN and CLN were measured. An additional 20 benchmark genes were identified as important by both RLN and CLN models, and 37 and 49 were specifically identified by RLN and CLN models, respectively. For example, *TIMING OF CAB EXPRESSION 1* (*TOC1*) was previously shown to control photoperiodic flowering response when RLN was used as a proxy for flowering time^27^. Consistent with this, *TOC1* only had importance ranks above 95^th^ percentile in G models for RLN. In our prediction models, *TERMINAL FLOWER 1* (*TFL1*) was only important for CLN prediction, which is consistent with the previous finding that *tfl1* mutants showed significantly decreased CLN, but not RLN or days to bolting (another proxy for flowering time), compared with wild type (WT)^28^. Taken together, these results indicate that different omics datasets, ways to represent data, importance measures, environmental factors, and ways to measure traits must be considered to better capture the genetic basis of Arabidopsis flowering time and likely other complex traits.

### Identification of novel genes involved in regulating flowering time

To determine whether all the features relevant to benchmark flowering time genes are sufficient to predict flowering time, we built RF models using G, T and gbM features for 426 benchmark genes separately or combined (hereafter referred to as benchmark gene-based models). Compared with the corresponding full models built using the features for all genes, the benchmark gene-based models had significantly lower performance (**Fig. 5a**), suggesting that genes in addition to the 426 benchmark genes are involved in regulating flowering time. To test this, we selected the 426 most important genes that were not benchmark genes (hereafter referred to as “novel” genes) from the full model, which was built using G/T/gbM features combined for all genes (**Table S17**). We found that the novel gene-based gbM model (built using the gbM features of these novel genes) performed significantly better than the benchmark gene-based gbM model, but not the corresponding G and T models (**Fig. 5a**). This may be attributed to the fact that most benchmark flowering time genes were identified using genetic approaches (e.g., via forward genetic screens or GWAS) and/or transcriptomic data (e.g., via gene differential expression analysis), rather than methylomic data.

**Fig. 5.**
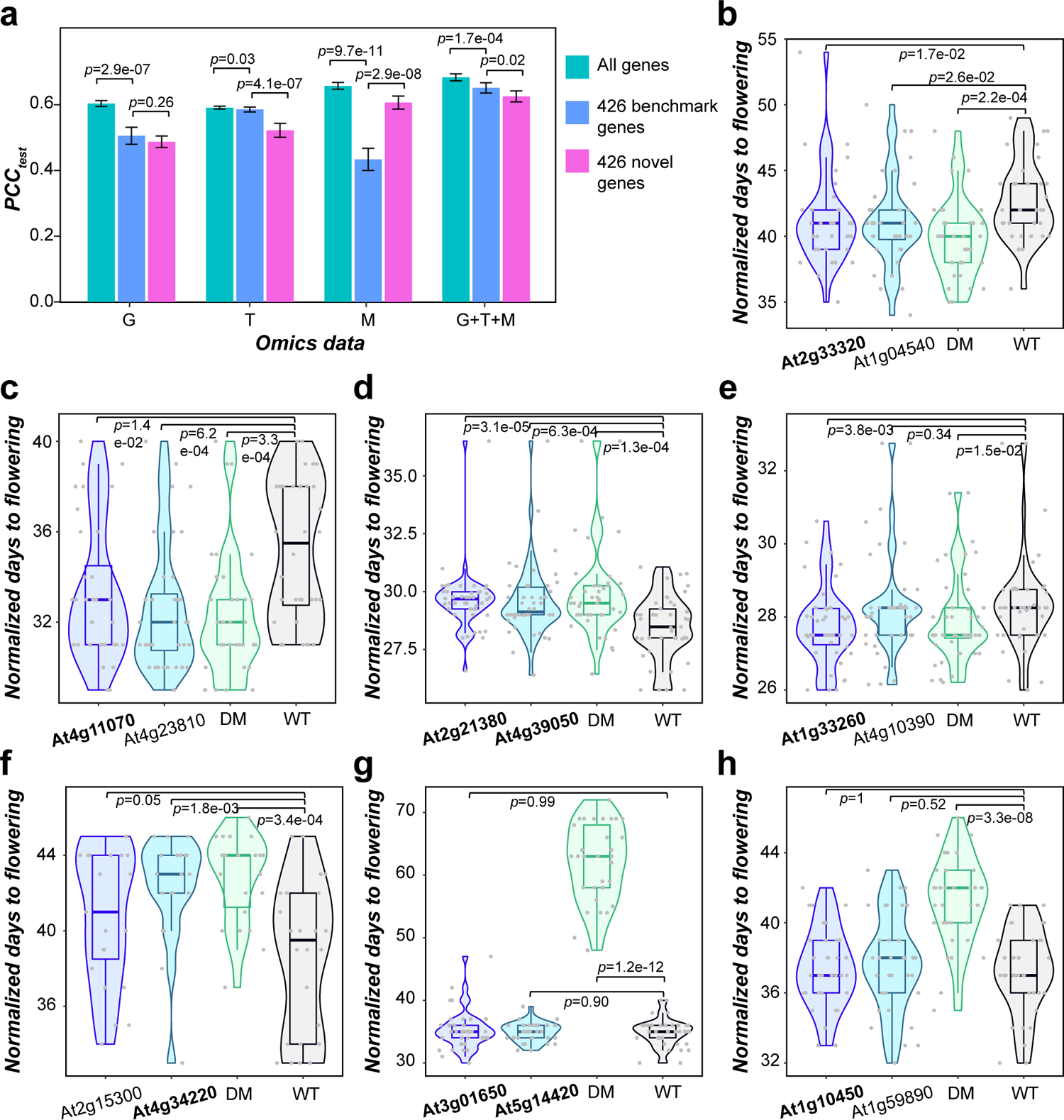
Identification of novel genes involved in flowering time regulation. (**a**) Accuracy of RF flowering time prediction models using all features (green), only features related to benchmark flowering time genes (blue), or only features related to the top 426 novel genes (magenta). G, T, and M: genomic, transcriptomic, and methylomic data, respectively. *p*-values are from Student’s *t*-test. (**b-h**) Statistical analysis of flowering time in single-gene mutants of important genes (bold font) and their paralogs (regular font), double mutant (DM) with loss of function of both paralogs, and wild-type (WT) plants. *p*-values are from two-sided Wilcoxon Rank Sum tests. Flowering time was normalized across flats within blocks (see **Methods**). Horizontal line in the box: median value; box range: interquartile range (IQR), 25^th^ (Q1) to 75^th^ percentile (Q3); whisker below box: Q1–1.5 IQR to Q1; whisker above box: Q3 to Q3+1.5 IQR; violin plot: distribution of datapoint values; dot: datapoint from an individual plant.

To validate the functions of important genes in flowering time, we took advantage of an existing dataset in our lab to compare flowering time in mutants of 21 non-benchmark genes that were identified as important and the WT (**Methods**, **Table S18**,**19**). Six of these 21 genes affected flowering time when mutated (**Table S18**; **Fig. 5b-h**). For example, a loss-of-function mutant of *AT4G11070* (*WRKY41*), which controls seed dormancy^29^, flowered significantly earlier than WT (**Fig. 5c**). Consistent with this, *WRKY41*’s homolog *Dlf1* regulates flowering in rice^30^. Another three genes had loss-of-function effects on flowering when mutated along with their paralogs (**Fig. 5g,h**, **Table S18**). The remaining 12 genes did not have a significant effect on flowering time when mutated, alone or with their paralogs. One potential explanation is that our validation experiments were conducted in the Col-0 genetic background, but the important genes were identified in models built across multiple accessions.

To check whether our models perform well in identifying genes that function in flowering time, we also measured the flowering time of the loss-of-function mutants of 37 “non-important” (importance rank ≤95^th^ percentile) genes and their paralogs. We found that 43.2% (16 genes) had significantly altered flowering time when mutated (**Table S18,19**). This percentage is only slightly lower than that of experimentally validated important genes (52.2%). One potential explanation for the similar percentages of predicted important and “non-important” genes with experimentally validated roles in flowering time is that, as discussed above, the importance of these genes may be dependent on the accessions and/or environmental conditions. In addition, there are far more features (e.g., >20,000 T features) than instances (only 383 accessions) in our models. Therefore, we do not have enough power to detect variants with significant but lower degrees of contribution to flowering time. Our false negative rate when using importance rank as a criterion is expected to be high.

### Accession-dependent contributions of genes to flowering time prediction

Thus far, we have evaluated the contribution of G, T, and gbM features associated with genes to the prediction of flowering time across accessions by dissecting the models through global interpretation^31^. However, some genes may be important contributors to flowering time only in specific accessions. To assess this, we determined the contributions of important features to flowering time in each accession (local interpretation) by examining the SHAP value for each feature in each accession (individual SHAP values, as opposed to the averaged value discussed earlier, see **Methods**). Here, a positive SHAP value for a feature in an accession means that the value of the feature in that accession contributed to a higher predicted trait value, i.e., longer flowering time. A negative SHAP indicates the opposite: the feature value contributed to reduced flowering time in an accession. The absolute SHAP value describes the degree of feature contribution to trait prediction. Here we first present the SHAP values of the top 20 important genes from the T model in detail as an example because more benchmark genes were among the top 20 genes in this model (**Fig. S7a**) than in the G (**Fig. S7b**) and gbM models (**Fig. S7c**). Organizing the accessions into clusters based on SHAP values of the top 20 T features allowed us to examine the way that different features contribute to flowering time prediction. The accessions we examined formed eight clusters (**Fig. 6a**), and flowering time varied greatly across these clusters (**Fig. 6b**).

**Fig. 6.**
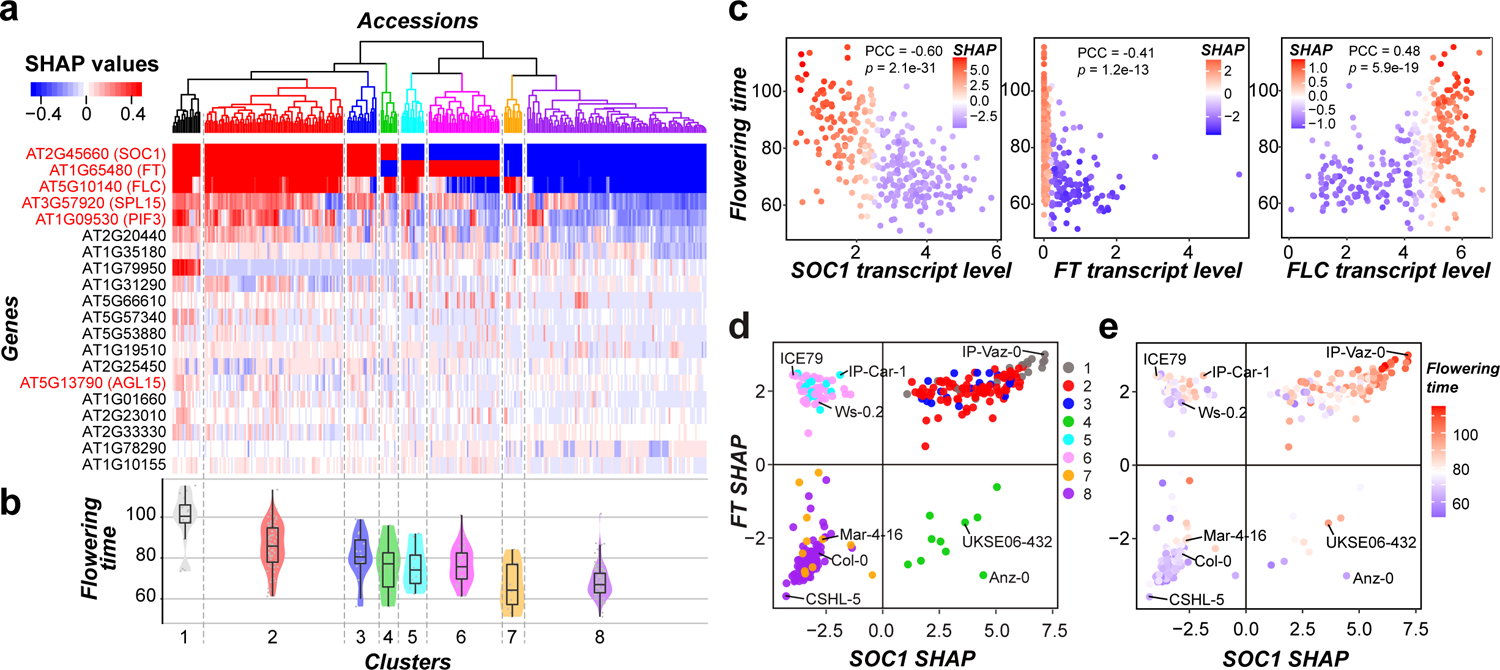
Accession-dependent effects of transcriptome features on flowering time. (**a**) Heatmap shows the SHAP values of the top 20 genes (y-axis) for each accession (x-axis). Benchmark gene names are in red on the left of the heatmap. (**b**) Violin plot shows the distribution of flowering time of accessions within each cluster. Horizontal line in the boxplot: median value; box range: interquartile range; dot: individual accession. (**c**) Correlation between *SOC1* (left), *FT* (middle), and *FLC* (right) expression levels with flowering time. Color scale: SHAP values for *SOC1*, *FT*, and *FLC* when using expression to predict flowering time. PCC: Pearson correlation coefficient, showing correlation between gene expression (x-axis, log[TPM + 1]) and flowering time (y-axis). (**d-e**) Relationships between *SOC1* and *FT* SHAP values among accessions; example accessions are labeled. Colors in (**d**): membership of accessions in clusters specified in **(a)**; color scale in (**e**): flowering time.

In cluster 1 and 2 accessions (**Fig. 6a,b**), the SHAP values for *SOC1*, *FT*, and *FLC* expression were all positive, indicating they contribute to longer flowering time. Their positive contribution is mainly due to lower *SOC1* and *FT* expression and higher *FLC* expression compared with other accessions (**Fig. 6c**). In cluster 8 accessions (**Fig. 6a,b**), all three genes have negative SHAPs, and the shorter flowering time is due to higher *SOC1* and *FT* expression but lower *FLC* expression. This is consistent with earlier findings that *SOC1* and *FT* promote flowering^32^, *FLC* represses flowering^33^, and the expression levels of *SOC1* and *FT* are negatively correlated with *FLC* expression^34^. This coupling between *SOC1, FT* and *FLC* in terms of expression (**Fig. 6c, Fig. S8a,b,f,g**), SHAP values (**Fig. 6d, Fig. S8c**), and flowering time contribution (**Fig. 6e, Fig. S8e**) is the predominant flowering time regulatory mechanism for most accessions. However, the action of these components can be decoupled. For example, in cluster 4, 5, and 6 accessions (green, cyan, and magenta; **Fig. 6d**), the SHAP values of *SOC1* and *FT* have opposite signs, which is associated with moderate flowering time values (**Fig. 6b,e**). Another example is cluster 7 (orange, **Fig. 6d**) where, despite the positive contribution of *FLC* (**Fig. S8c**), both *SOC1* and *FT* contribute negatively to flowering time (**Fig. 6d**). This coupling and uncoupling between *SOC1*, *FT*, and *FLC* expression across accessions indicates that regulation of flowering time may be more complex than what has been reported. Accession-dependent contributions of other important T features are also observed, although to a much lower extent. An example is *SQUAMOSA PROMOTER BINDING PROTEIN-LIKE 15* (*SPL15*); clusters 2 and 4–8 can be divided into sub-clusters depending on whether *SPL15* expression contributes positively to flowering time in an accession (**Fig. 6a**).

We also clustered accessions using SHAP values of the top 20 important features from G and gbM models, and obtained eight and nine clusters, respectively (**Fig. S9a,c**), which resembled the distribution of T model-based clusters in that the clusters could be characterized by a few top features. For example, the second (*AT4G38550*) and the third (*AT1G35610*) most important genes from the gbM model can be coupled or uncoupled with *FLC* in a similar fashion as *SOC1* and *FT* (**Fig. S9c**). The different omics data types yielded clusters with different accessions (**Fig. S10**). These findings further demonstrate that different omics data reveal different contributors to flowering time variation among accessions, and that the different effects of genes on flowering time among accessions can be disentangled through model interpretation. In addition, knowledge about flowering regulation in some accessions may not be generalizable to others, as demonstrated by the identification of different benchmark genes when different sets of accessions were used (**Table S10,S12**). This may be explained by the differences in genetic backgrounds among accessions^25^ and the high complexity of genetic interaction networks regulating flowering^35^. The accession-dependent effects of genes on flowering may also partially explain the low degree of enrichment of benchmark genes among important genes in our original G, T, and gbM models because these benchmark genes were predominantly discovered in the Col-0 accession.

### Genetic interactions revealed through integration of multi-omics data

In the above sections we showed that different types of omics data revealed overlapping but mostly distinct genes impacting flowering time. Next, we asked whether combining different types of omics data improves the prediction accuracy. We found that combining G and T or all three datasets improved model performance for RF models, but not for rrBLUP models (**Fig. 7a**). Because the RF algorithm considers non-linear feature combinations while rrBLUP does not, the better RF model performance suggests that the inclusion of interactions between features from different omics data may have improved model performance. To evaluate this possibility, we established an additional RF model, which only included G+T+gbM features relevant to the 378 benchmark flowering time genes (see **Methods**) to facilitate model interpretation. We used the SHAP approach^36^ to identify feature interactions (see **Methods**), where the contribution of feature X to trait prediction is influenced by values of feature Y. The SHAP feature interaction can help us identify potential genetic interactions between genes or variants, such as epistasis^37^. We identified 7,186 feature interactions, including all six possible combinations between omics data types: G-G, T-T, gbM-gbM, G-T, G-gbM, and T-gbM (**Table S13**). G-T, T-gbM, and T-T interactions were the most prevalent (**Fig. 7b**). The three most important genes in the T model—*SOC1*, *FT*, and *FLC* (**Fig. 6a**)—had the highest numbers of feature interactions (752, 658, and 287, respectively), consistent with their reported functions as floral integrators^32^, which receive floral promotion or inhibitory signaling inputs from distinct pathways (**Fig. 7c**, **Table S20**).

**Fig. 7.**
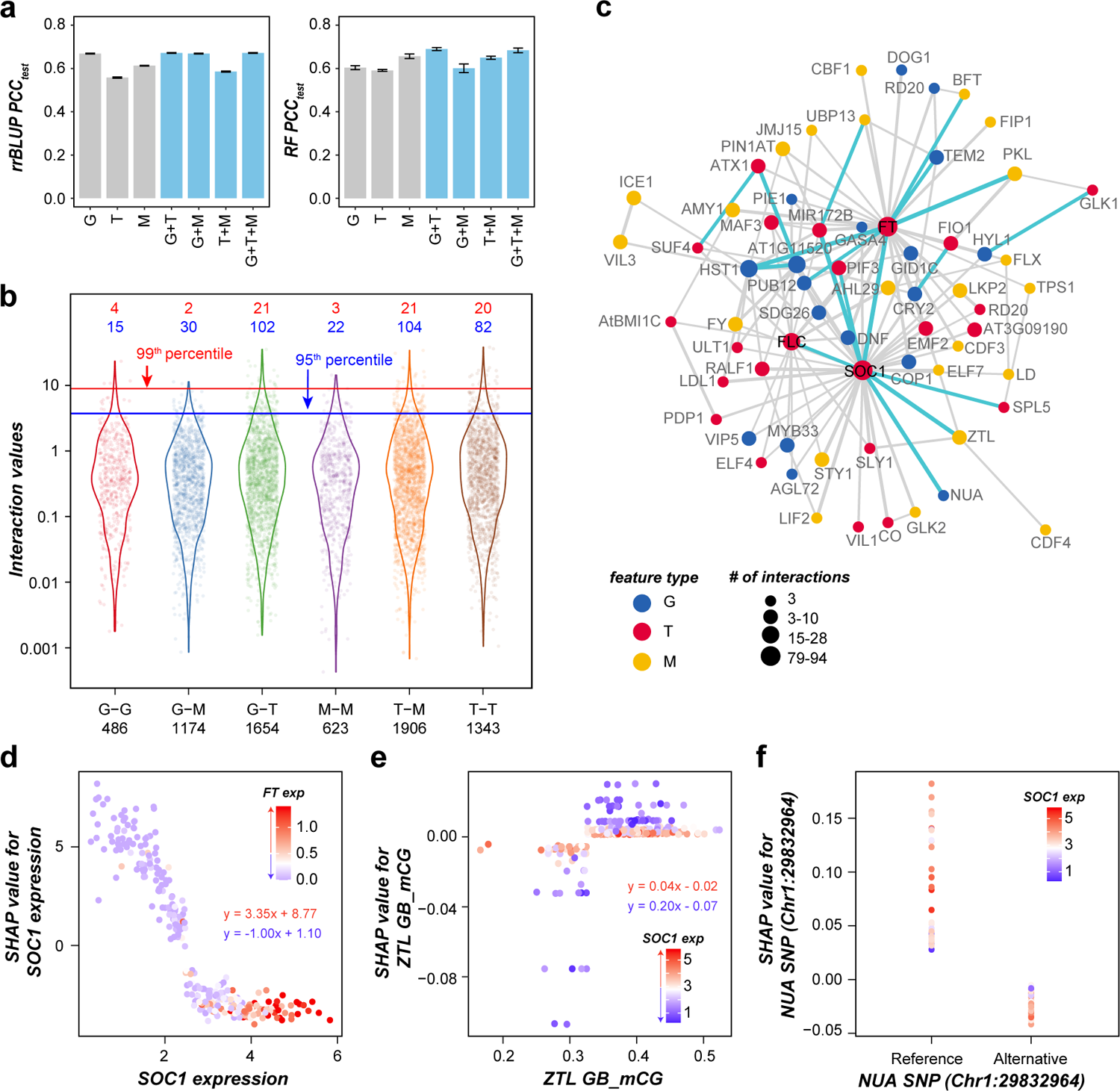
Prediction models integrating all omics data types and putative interactions between flowering time genes. (**a**) Prediction accuracy of flowering time for models built using single (gray) and combined (blue) omics data. Left panel: rrBLUP models, right panel: RF models; G, T, and M: genomic, transcriptomic, and methylomic data, respectively; error bars: standard deviation of 10 runs. (**b**) Distributions of SHAP interaction values for different types of feature interactions. X-axis label: interaction type and number of identified interactions. Red and blue dashed lines: 99^th^ and 95^th^ percentiles of interaction values, respectively; numbers in red and blue font: numbers of feature interactions above the 99^th^ and 95^th^ percentiles, respectively, for each type of interaction. (**c**) Network of features with ≥ 3 interaction values above the 95^th^ percentile. Blue, red, and orange points: G, T, and gbM features, respectively; node size: number of interactions; edge thickness: log_10_(interaction value + 1)/2; blue edges: 18 interactions that ranked above 20^th^ and had nodes with size ≥ 3; gray edges: all the other interactions; black font: three genes with the highest number of interactions. (**d-f**) Interactions between *SOC1* expression and *FT* expression (**d**), *ZTL* CG-type gene-body methylation (**e**), and a *NUA* SNP (**f**). Equations in (**d,e**) are for regression of the given feature values (x-axis) and the corresponding SHAP values (y-axis) in accessions with certain *FT* (**d**) and *SOC1* (**e**) expression values (log[TPM + 1]): red font: *FT* expression ≥ 0.5 or *SOC1* expression ≥ 3; blue font: *FT* < 0.5 or *SOC1* < 3. TPM: transcripts per million.

Among the top 20 interactions with the highest interaction values, five were T-T interactions of *SOC1* with *FT* (ranked 1^st^), MIR172B (2^nd^), *SPL5* (13^th^), *FLC* (19^th^), and *PHYTOCHROME INTERACTING FACTOR 3* (*PIF3*) (20^th^) (**Fig. 7c,d**, **Fig. S11a-e**). SOC1 was previously shown to regulate *MIR172B*^38^ and *SPL5*^39^ by directly binding to their promoters. However, no direct biological interaction between *SOC1* and *PIF3* has been reported. Also among the top 20 interactions, two were T-gbM and T-G interactions of *SOC1* with *ZEITLUPE* (*ZTL*, ranked 5^th^, **Fig. 7e**) and *Nuclear Pore Anchor* (*NUA*, ranked 6^th^, **Fig. 7f**). *ZTL* and its two homologs, *LOV KELCH PROTEIN 2* (gbM-T interaction with *SOC1* ranked 121^st^, **Fig. S11f**) and *FLAVIN-BINDING, KELCH REPEAT, F-BOX 1* (gbM-T interaction with *SOC1* ranked 3333^rd^, **Fig. S11g**), function as E3 ubiquitin ligases and regulate flowering time regulators, such as CONSTANS (CO) and FT^40^ No direct biological interaction has been reported between *SOC1* and *ZTL* or its two homologs, whereas the interaction between *SOC1* and *NUA* was reported before: expression of *SOC1* was higher in *nua-1/4* mutants compared with WT^41^. In addition to these T-T, T-M, and T-G interactions, we also observed G-G, G-gbM, and gbM-gbM interactions that have not been reported before: a G-G interaction between *AT1G11520* and *HASTY 1* (*HST1*) ranked 8^th^ (**Fig. S11h**), gbM-gbM interaction between *VIN3-LIKE 3* (*VIL3*) and *INDUCER OF CBF EXPRESSION 1* (*ICE1*) ranked 25^th^ (**Fig. S11i**), and G-gbM interaction between *AT1G11520* and *FY* ranked 45^th^ (**Fig. S11j**). These novel interactions might indicate potential biological interactions between genes.

Furthermore, by examining the feature interactions in detail, we found some interesting patterns. For example, the T-gbM interaction between *SOC1* and *ZTL* illustrates how *SOC1* and *ZTL* interact across accessions (**Fig. 7e**): in accessions with higher *SOC1* expression, contributions of *ZTL* CG-type gene-body methylation were near-zero regardless of the methylation levels of *ZTL*, whereas in accessions with lower *SOC1* expression, the contributions were larger either positively or negatively. A similar pattern was also observed for other interactions, such as a T-T interaction between *SOC1* and *MIR172B* (**Fig. S11b**). Taken together, these findings show that potential interactions at different molecular levels can be identified on a large scale by interpreting computational models integrating multiple omics data.

## Conclusion

We investigated the utility of G, T, and M data in predicting complex traits in Arabidopsis, and found that models built using these data types separately had similar performances. The flowering time prediction models built using different omics data identified different benchmark flowering time genes and other genes not previously reported to be involved in flowering. Even models built using different forms of a single type of omics data—M data—identified different sets of flowering genes. These results highlight the necessity of exploring different types of omics data when predicting complex traits. Even though there was only one time point for transcriptome data (rosette leaves right before bolting)^9^, key regulators of flowering time, such as *FLC*, *FT*, *SOC1*, and *SPL15*, were still identified in T-based rrBLUP and RF models. Considering that rrBLUP is an algorithm based on a linear model, this finding indicates that simple, linear combinations of the steady-state expression levels of these genes, which explain a significant portion of the flowering time variation among accessions, can be identified.

We identified important genes including the most well known flowering time genes; however, the false positive and false negative rates based on the benchmarks were high. Model performance and candidate gene identification can be further improved by incorporating substantially more accessions, using additional approaches to select, combine, and represent features, and incorporating data from more than one environment. In addition, the Arabidopsis T and M data used in this study were obtained from mixed rosette leaves harvested just before bolting, which poses two potential problems for complex trait prediction: 1) gene expression and methylation can be cell-type specific, thus noise in T and M data is inevitable due to cell heterogeneity; 2) the complex traits to be predicted may be specific to certain tissues, organs, or development stages; thus, T and M variation from leaves may not reflect trait variation in other contexts. The development of single-cell and spatial omics techniques, which allow the cellular landscape of epigenomes within a tissue to be measured, is expected to improve the prediction accuracy of complex traits. In addition, omics data in addition to G, T, and M data can be used to predict complex traits; these data include chromatin architecture, chromatin accessibility, and histone modification, changes of which have been shown to regulate flowering in Arabidopsis^39,42–44^.

That the effects of genes or variants on flowering differ among accessions is known in specific cases; for example, DNA demethylation has different effects on flowering in C24 and Landsberg *erecta*^45^, and some accessions carry non-functional alleles of the *FRI* gene^25^. Here, we show that accession-specific effects of different flowering time genes can be revealed by interpreting machine learning models. The outcome also revealed the accession-dependent effects that were not documented previously. Interpretation of a non-linear model (RF) also allowed the identification of known and novel interactions among G, T, and gbM features of benchmark genes. The novel interactions represent new hypotheses that require further experimental verification and are expected to provide insights into how these novel components and interactions contribute to the genetic basis of flowering time and potentially other complex traits.

## Methods

### Data preprocessing

Six traits for each Arabidopsis accession (**Table S1**) were obtained from two publications: (1) flowering time at 10 °C or 16 °C, which was scored as days until the first flower was open, was from^10^; (2) cauline leaf number (CLN), (3) rosette leaf number (RLN), (4) rosette branch number (RBN), (5) diameter of rosette (end point after flowering, DoR), and (6) length of main flowering stem (stem length, SL) were from^11^. For each trait, the pairwise Euclidean distances of trait values between accessions were calculated using the R package “rdist”. The Euclidean distances were first normalized between 0 and 1, and then the correlation or similarity of phenotypic trait values (pCor) among accessions was calculated as 1 - normalized Euclidean distance.

The genomic matrix (G) was downloaded from the 1001 Genomes database^10^ (http://1001genomes.org/data/GMI-MPI/releases/v3.1/SNP_matrix_imputed_hdf5/). This matrix only contains biallelic SNPs. SNPs with minor allele frequency < 0.05 were removed. For each SNP, the major allele was encoded as 1, and the minor allele was encoded as −1. The kinship matrix was generated from G using the KinshipPlugin with the centered Identity By State method^46^ implemented in TASSEL v5^47^. The kinship typically refers to the degree of genetic relatedness between accessions, and was used here as the proxy for the similarity of G among accessions. The first five principal components (used as the proxy for population structure) were generated using the PrincipalComponentsPlugin implemented in TASSEL v5.

For the transcriptomic (T) data^9^, the normalized read counts were downloaded from NCBI (https://www.ncbi.nlm.nih.gov/geo/download/?acc=GSE80744), and were used to calculate the transcripts per million (TPM) using the function “calculateTPM” from the R package “scater”^48^. The TPM values were then transformed to ln(TPM+1). The transformed TPM values were used to calculate the expression correlations (eCor) among accessions, and the resultant Pearson correlation coefficient (PCC) values were normalized between 0 and 1 to make them comparable with pCor, which ranges from 0 to 1.

For the methylomic (M) data^9^, the gene-body methylation values (the number of reads with methylated cytosines in a gene body divided by the total number of reads with both cytosines and thymines) were downloaded from http://signal-genet.salk.edu/1001.php. For accessions with multiple replicates, the median values were calculated across replicates. The missing values were imputed using the k-Nearest Neighbors approach (“KNNImputer” function in sklearn.impute)^49^ as follows: missing values in the training subset (for the data split to training and test subsets, see the Methods section “**Predictive modeling**”) were first imputed, then the imputation was transformed to the test subset. To mitigate potential influences of the selection of k on the imputed missing values, the imputation was conducted five times with different k values (3, 4, 5, 6, and 7), and the means of imputed values were used. The imputed gene-body methylation values were used to calculate the gene-body methylation correlation (mCor) among accessions, and the resultant PCC values were normalized between 0 and 1.

### Additional formats of methylomic data

The frequency of cytosine sites can vary within a genic region, and the methylation level at each cytosine site (i.e., proportion of mapped reads with methylated cytosine) can also vary because of cell heterogeneity; therefore, the gene-body methylation obscures information such as heterogeneity in methylation levels over the length of a gene. To overcome these shortcomings, we used six more types of M data (**Fig. S3**) to represent single site-based methylation (ssM) for predicting flowering time: (1) methylation status of single cytosine sites (i.e., 1, indicating methylated, or 0, indicating not methylated, hereafter referred to as presence/absence [P/A] of methylation) across the whole genome; (2) methylation proportion (i.e., proportion [Prop] of reads that were methylated at that site) for single cytosine sites across the whole genome; (3,4) methylation profiles along each gene, i.e., the methylation status (3, mean_P/A) or proportion (4, med_Prop) across upstream, gene-body, and downstream regions divided into 30 bins; (5,6) clusters of genes with similar methylation profiles when methylation status (5, P/A_clu) or proportion (6, Prop_clu) was considered.

The methylation base calls for each reference site^9^ were downloaded from NCBI (accession ID GSE43857). Missing values in the single cytosine site methylation matrix were treated as follows: (1) if the reference site was cytosine and there was a SNP for this site in accession X, or if the reference site was not a cytosine and this site was not a SNP for X, the methylation call for this site in X was encoded as 0; (2) if the site’s base in accession X was unknown, or the site in X was a cytosine, but there were no reads mapped to this site, then the values were marked as missing to be imputed in the same way as for the gene-body methylation matrices, except that for the P/A of methylation for a single site, the final imputed value was rounded to either 0 or 1. After imputation, sites with the same value (either 0 or 1) in > 95% accessions (similar to 5% minor allele frequency) were removed. To balance the trade-off between the number of methylation sites included in the analysis and the number of missing data points, we first ordered the sites according to the proportion of missing values across accessions. Three thresholds were explored: 90^th^, 75^th^, and 50^th^ percentile (the corresponding datasets are referred to hereafter as 90per, 75per, and 50per, respectively), sites above which had missing values in ≤ 2, 8, and 38 accessions, respectively (**Fig. S12a**). We found that the dataset 50per tended to have the highest performance in rrBLUP models while dataset 90per led to the highest performance in RF models (**Fig. S12b,c**). To include more single-site methylation information, matrices using the 50^th^ percentile as a threshold were used for subsequent analysis.

To bin the methylation levels of single sites for gene-based regions, the gene regions were split into 30 bins as shown in **Fig. S12d**. For each bin, a summary statistic representing the methylation data was calculated: for the methylation P/A, this statistic is the mean number of methylated cytosines in the bin; for methylation proportion, this statistic is the median of the methylation proportion values (a number between 0 and 1) in a bin. To summarize the methylation profiles of genes across accessions, the binned methylation data were formatted into vectors, with one vector of length 30 (the number of bins) for each gene in each accession. These vectors were clustered using K-means clustering with 30 clusters. For each gene in each accession, 30 new features (i.e., whether the vector of the gene [methylation profile] belongs to each of the 30 clusters, 1 for the cluster the gene belongs to, 0 for all the other clusters) were produced. Finally, for each feature matrix, columns containing only zeros were removed. These processes were performed on each methylation type separately (CG, CHG, CHH), and matrices of three methylation types were combined together, resulting in 16 final feature matrices (16 “binned” columns in **Table S3**). To simplify our story, these 16 matrics were only used to establish predictive models for flowering time.

### Predictive modeling

Twenty percent of the accessions were randomly held out as the test set, which was used to evaluate the performance of final models and was never used in the model training. The remaining 80% of accessions were used to train the models. A five-fold cross-validation (CV) scheme was conducted in the model training. First, the remaining 80% of accessions were randomly split into five folds. Next, accessions in four folds (referred to as the training set) were used to build the model, and accessions in the fifth fold (validation set) were used to evaluate the model performance. Finally, this training-validation step was conducted five times to make sure each fold was used as a validation set once. This five-fold CV scheme was repeated 10 times. The PCC was calculated between true trait values and the predicted values, and the average PCC among the 10 runs was used to measure the model performance on the CV set.

The R package “rrBLUP” (ridge regression Best Linear Unbiased Prediction)^12^ and the scikit-learn class “RandomForestRegressor”^13,49^ were used to build the predictive models. rrBLUP is a commonly used genomic prediction approach with mixed models, and the key function used in this study is the “mixed.solve”, with the equation: y = μ + Xg + e, where y is the vector of trait values, μ is the overall mean of trait values for the training set, X is the feature matrix, g is the feature effects matrix (or coefficient matrix), and e is the vector of residual effects. The coefficients of features were estimated within each fold of CV and were used to predict trait values for accessions in the validation and test set. After five folds of CV, the prediction accuracies on the test set were averaged to evaluate the model performance, and the coefficients were also averaged.

RandomForestRegressor is a meta estimator that builds various regression decision trees using a number of sub-samples of the dataset. The resulting trees are then aggregated through averaging into a single ensemble model. GridSearch was used for hyperparameter tuning, where spaces of the parameters -max_depth and -max_features were [3, 5, 10] and [0.1, 0.25, 0.5, 0.75, “sqrt”, “log2”, “None”], respectively. The parameter -n_estimators was set as 100. The best parameter combination was selected according to the model performances in CV. A final model was built using all the accessions (including accessions in both the training and validation sets) with the best parameter combination, and was applied on the test set to evaluate the model performance. To establish the baseline for the genomic prediction, we built predictive models based simply on population structure (defined as first five principal components from genetic markers, as mentioned above, blue dotted line in **Fig. 2b**) using the training subset. Models based on individual omics data outperformed those based on population structure for flowering time, RLN, and CLN, but not for DoR, RBN, and SL. In addition, prediction performances for models based on individual omics data and population structure on the CV and test sets for DoR differed dramatically in both rrBLUP and RF models (**Fig. 2b**, **Fig. S2**), which is indicative of high heterogeneity in DoR among accessions; the accessions in the training and test sets might have different genetic features affecting DoR. The prediction performances for RBN and SL were relatively low (< 0.2 and ∼0, respectively, **Fig. 2b**, **Fig. S2a**) no matter which omics data or population structure was used to build the models. Therefore, variation in RBN and SL might be mainly explained by the environment and/or genotype-by-environment interactions.

To remove the potential confounding effects of kinship on mCor, we estimated linear regression models between mCor and kinship for training accessions (i.e., matrices containing only mCor and kinship values between all accessions (in both training and test sets) with accessions in the training set using the “lm” function in R, and then used the residuals of mCor to build a new Random Forest (RF) model.

### Feature importance

Three measures were used to evaluate the importance or contributions of features to the prediction of complex traits: (1) RF “gini” importance, which reflects the impurity decrease when a feature is fed to the model^50^; (2) coefficients of features in the rrBLUP regression models; and (3) SHapley Additive exPlanations (SHAP) values, which reflect the contribution of a feature to the prediction of a complex trait^18^. The average absolute SHAP value of a feature for all the instances (local feature importance) was used to measure the global contribution of the feature to model prediction (global feature importance). The former two measures interpret global feature contributions to the model predictions and tend to assign non-zero importance values to all or most features (**Fig. 3, Fig. S4a-c**). In contrast, SHAP values provide local interpretations and tend to assign zero for features that have no contribution to the prediction of flowering time for individual instances (i.e., accessions in this study, **Fig. S4d-f**). A positive SHAP value indicates an instance is predicted to have a higher trait value with a given feature than when that feature is removed from the model, and vice versa. The higher the absolute SHAP value, the more a feature contributes to trait prediction for the instance in question.

### Benchmark flowering time genes

We downloaded 378 and 48 benchmark flowering time genes (**Table S2**) from the FLOR-ID database (http://www.phytosystems.ulg.ac.be/florid/)^15^ and TAIR (https://www.arabidopsis.org), respectively; these genes are known to be involved in flowering in Arabidopsis. To compare the importance of flowering genes across models built using different omics data, the highest importance (i.e., absolute coefficient in rrBLUP models, RF feature importance, and absolute SHAP values) of all SNPs (or methylation sites) within a gene was used as the gene-based importance. When all three types of gene-body methylation data (i.e., CG, CGH, and CHH) were used together, the highest feature importance of the three types was used as the gene-based importance.

To understand why no more benchmark genes at significant levels were identified than random chance for most combinations of omics datasets and importance measures, we explored the following three potential reasons. First, we determined whether the threshold affects the number of identified benchmark genes by increasing the threshold of gene importance scores to the 99^th^ percentile. The higher the threshold, the fewer benchmark genes were identified, but also the fewer genes were expected (1% for threshold at 99^th^ percentile). We found that there were significantly more benchmark genes identified than random (1%) only when SHAP values from models built using G and T data were used to identify important genes (**Table S11**), indicating that SHAP values are able to reveal the most important genes (99^th^ vs 95^th^ percentile) for flowering time prediction. However, for all the other combinations of datasets and feature importance measures, no significant differences were observed between results when the 95^th^ and 99^th^ percentiles were used as thresholds, indicating that the choice of threshold only had a minor effect on the identification of benchmark genes.

Second, we asked whether the sample size of accessions (306 accessions in the training set, for which data for six traits and all the G, T, and M data were available) was too small. To test this, we repeated the analysis using all 618 accessions (494 accessions were used to train the model, for which all the G, T, and M data and flowering time information were available). Generally, no more benchmark genes were identified than when 306 accessions were used to train the models (**Table S12**), suggesting that decreasing the number of accessions included in the model from 618 to 383 was not a major factor affecting the identification of benchmark genes.

Third, since the functions of benchmark genes in flowering may have been determined under different conditions (e.g., under a different temperature or photoperiod), we investigated whether the use of flowering time data measured under only one condition (at 10 °C)^10^ explained the failure to identify more benchmark genes than expected. We reasoned that benchmark genes showing effects on flowering under multiple conditions when mutated or overexpressed would be more likely to contribute to flowering at 10 °C than genes showing effects only under one condition. As expected, we found that genes contributing to flowering under two different conditions (short days [SDs] and long days [LDs], obtained from the FLOR-ID database; unfortunately there were no data for different temperatures) were more likely to be identified than those contributing to flowering only under one condition (**Table S10,S11**, **Fig. S13**). In addition, when predicting flowering time measured at 10 °C or 16 °C, genes had different contributions (**Table S14**). These findings suggest that the conditions under which the target traits were measured affect which genes are identified as important genes, consistent with the different QTLs identified for flowering time measured at different temperatures^10^.

In summary, the threshold used and the number of accessions used to train models only had minor, if any, effects on the number of known flowering genes identified. Nevertheless, our feature importance-based approaches outperformed the GWAS approach from a previous study^10^, where only five QTLs were identified. Thus, we continued with our original strategy, using the important genes with feature importance scores above the 95^th^ percentile from models built using 306 accessions, for subsequent analysis.

### Plant materials and assessment of flowering time

The potential function in flowering time of 21 genes with importance values above the 95^th^ percentile (for at least one model based on one of the three omics data combined with one of three importance measures) was validated experimentally. Flowering time data for T-DNA insertion mutants of these 21 important genes and another 37 non-important genes in the Arabidopsis Col-0 background were obtained from an existing dataset in our lab, which was generated to assess the degree of genetic redundancy between duplicate genes (**Table S18**). Generation and genotyping of mutants and the planting and growth conditions were described in detail previously^51^. Stock numbers for single mutants for these genes are listed in **Table S18**. The double-mutants were produced by crossing the corresponding single mutant lines. In brief, seeds for homozygous mutant and WT sibling plants (using seeds collected from two to six plants per genotype; referred to as sublines) were planted to compare the flowering time between mutants and WT.

Days to flowering (number of days from placing flats containing seeds in the growth chamber until the appearance of the first open flower) were recorded for at least 40 plants per genotype (**Table S19**). Within each block, days to flowering observed for plants grown in different flats was first normalized using the function “normalize” of the R package “broman”. Average differences in flowering time between mutants and WT were calculated, and the significance of differences was assessed using the two-sided Wilcoxon rank sum test.

### Feature integration

To investigate interactions among G, T, and gbM features of benchmark flowering time genes, we first built a model using all the features (G + T + gbM) related to the 378 benchmark genes. Then, to simplify the analysis, only one feature each for G (i.e., the SNP having the highest importance rank among other SNPs in a gene), T, and gbM (i.e., CG-, CHG-, or CHH-type gene-body methylation, having the highest importance rank among others in a gene) was kept for each gene. This resulted in 1227 features for the 426 genes. A new model was built using these top features, and the interactions among these features were calculated using the “TreeExplainer” and “shap_interaction_values” functions of the SHAP package^36^. The SHAP interaction of feature i and feature j can be interpreted as the difference in SHAP values of feature i with and without the feature j^36^. The higher the value, the stronger the interaction.

## Supplementary figure legends

**Fig. S1 Similarity of flowering time, kinship, eCor, and mCor among accessions.** Heatmaps showing phenotypic (pCor, of flowering time, **a**), genomic (kinship, **b**), transcriptomic (eCor, **c**), and gene body methylation (mCor, **d**) similarities among 383 accessions. The ordering of accessions is the same across **a**-**d**. Phenotypic similarities were measured as 1-euclidean distance. pCor, eCor, and mCor were normalized between 0 and 1. Color scales are shown above the heatmaps. Yellow boxes: clusters of accessions with relatively higher phenotypic similarities.

**Fig. S2 Model performance using three different types of omics data on test and cross-validation data.** (**a**) Performance on the test set using Random Forest (RF) models. The corresponding model using ridge regression Best Linear Unbiased Prediction (rrBLUP) is shown in **Fig. 2b**. (**b**,**c**) Model performance on the cross-validation (CV) set using rrBLUP (**b**) and RF (**c**) models. (**d**) Correlation between model performance (y-axis, left panel: rrBLUP models; right panel: RF models) and the correlation between the trait value and omics data similarity among accessions (x-axis) as shown in **Fig. 2a**. Each dot represents the model performance and correlation for a single trait and omics data type. (**e**) Correlation between the rrBLUP and RF model performances on the test set for different omics data types. PCC: Pearson correlation coefficient; G: genome; T: transcriptome; mGeneBody: gene-body methylation; K: kinship; eCor: expression correlation; mCor: methylation correlation.

**Fig. S3 Prediction performances for flowering time using different types of omics data.** Prediction performances for flowering time on the test (**a**,**c**) and validation (**b**,**d**) sets for rrBLUP (**a**,**b**) and RF (**c**,**d**) models. CV: cross-validation; PCC: Pearson correlation coefficient; G: genome; T: transcriptome; gbM: gene-body methylation; CG, CHG, CHH: three types of methylation; Single site: methylation level at a single cytosine site, where presence/absence (P/A) of methylation at a site was encoded as 1/0, and the degree of methylation at a site was defined as the proportion (Prop) of methylated reads over all the reads mapped to that cytosine site; Binned: the proportion of methylated cytosine sites over all cytosine sites within a bin (mean_P/A, for how the gene regions were binned, see **Fig. S12d**), or the median degree of methylation within a bin (med_Prop); P/A_Clu: methylation type membership of genes in clusters grouped based on the mean_P/A across all bins; Prop_Clu: methylation type membership of genes in clusters grouped based on the med_Prop across all bins.

**Fig. S4 Correlation of feature importance between different types of omics data.** Density scatter plots showing the correlation of the importance scores between genomic (G), transcriptomic (T), and methylomic (M) features of genes when rrBLUP coefficients (**a-c**) and SHapley Additive exPlanations (SHAP) values (**d-f**) were used as measures of importance. The absolute values of rrBLUP coefficients and SHAP values were used. All feature importance scores < 10^-6^ were changed to 10^-6^. Red dashed line: 95^th^ percentile of feature importance. Density plot color: density of genes. ρ: Spearman’s rank correlation coefficient. (**g**) Confounding effects of kinship on mCor when predicting complex traits. Purple and blue bars: performance on the cross-validation (CV) set for models built using mCor matrix and using residuals of mCor with confounding effects of kinship removed (see **Methods**), respectively; green and yellow bars: performance on the test set for models built using mCor matrix and using residuals of mCor, respectively. Error bars: standard deviation of 10 runs. Blue dashed line: performance of models built with first five PCAs of genomic variation (population structure) evaluated on the test set. *p*-values: Student’s *t*-test. RLN: rosette leaf number; CLN: cauline leaf number; DoR: diameter of rosette; RBN: rosette branch number; SL: stem length.

**Fig. S5 Relationship between different features of *CSTF77* and flowering time among accessions.** (**a,b**) Significantly different flowering times were observed between accessions with a methylated cytosine site (1) and accessions with a non-methylated cytosine site (0) at loci Chr1_6111193 (**a**) and Chr1_6113613 (**b**). (**c**,**d**) No significant differences in flowering time were observed between accessions with the alternative allele (1) and accessions with the reference allele (−1) at loci Chr1_6111716 (**c**) and Chr1_6111257 (**d**). *p*-values in (**a-d**) are from two-sided Wilcoxon Rank Sum tests. (**e-h**) Correlation between flowering time and *CSTF77* expression (**e**) and CG-(**f**), CHG-(**g**), and CHH-type (**h**) gene-body methylation among accessions. PCC: Pearson correlation coefficient.

**Fig. S6. Effects of different environmental conditions on gene importance for flowering time prediction.** (**a**) Different SHAP values indicating importance of gene expression for flowering time prediction for plants grown at 10 °C (x-axis) and 16 °C (y-axis). All the zero values were assigned to be 0.001. (**b**) Flowering time differences in accessions with non-functional (blue) and functional (red) *FRIGIDA* (*FRI*) alleles at 10 °C and 16 °C. Horizontal line in the box: median value; box range: interquartile range (IQR), 25^th^ (Q1) to 75^th^ percentile (Q3); whisker below box: Q1–1.5 IQR to Q1; whisker above box: Q3 to Q3+1.5 IQR; dots: outliers. (**c,d**) rrBLUP coefficient (**c**) and RF gini importance (**d**) values of T features for flowering time prediction for plants grown at 10 °C (x-axis) and 16 °C (y-axis). In (**d**), zero values were assigned to be 1e-9.

**Fig. S7 Top features with the highest SHAP values when three types of omics data were used.** SHAP values of the top 20 genes corresponding to transcriptomic (**a**), genomic, (**b**) and gene-body methylation (**c**) features. Left panel shows original SHAP values; in the right panel the SHAP values are extended to the same scale for the sake of comparison. Color scale in (**a,c**): feature values. In (**b**), red color: alternative allele, blue color: reference allele. Features related to benchmark flowering time genes are highlighted in red font.

Fig. S8 Correlation of SHAP values for *SUPPRESSOR OF OVEREXPRESSION OF CO 1* (*SOC1*), *FLOWERING LOCUS T* (*FT*), and *FLOWERING LOCUS C* (*FLC*) expression with flowering regulation. (**a,b**) Correlation between *SOC1* and *FT* SHAP values. Color scale in (**a**) indicates *SOC1* expression levels, and color scale in (**b**) indicates *FT* expression. (**c-g**) Relationships between *SOC1* and *FLC* SHAP values among accessions. Color in **c** indicates membership of accessions in clusters shown in **Fig. 6**. Clusters are based on SHAP values of the top 20 genes from flowering time prediction models using T data. Colors in **e**, **f,** and **g** indicate flowering time, *SOC1* expression, and *FLC* expression, respectively. Example accessions are indicated in (**a-g**).

**Fig. S9 Flowering time for accessions clustered based on SHAP values of the top 20 features from G and gbM data-based prediction models.** (**a**,**c**) Color in heatmaps shows the SHAP value of a SNP (**a**) or gene-body methylation level (**c**) (y-axis) for a given accession (x-axis). Features related to benchmark genes are highlighted in red font on the left of the heatmap. (**b**,**d**) Violin plots in **b** and **d** show the distribution of flowering time values for accessions within each cluster shown in **a** and **c**, respectively. Horizontal line in the box: median value; box range: interquartile range (IQR), 25^th^ (Q1) to 75^th^ percentile (Q3); whisker below box: Q1–1.5 IQR to Q1; whisker above box: Q3 to Q3+1.5 IQR; dot: individual accession.

**Fig. S10 Clusters of accessions based on SHAP values of features from G, T, and gbM data.** (**a-c**) The clusters of accessions in **a**, **b**, and **c** are the same as those in **Fig. S9a**, **Fig. 6a**, and **Fig. S9c**, respectively, except that only SHAP values for the top three features are shown. (**d**) Colors indicate the cluster memberships of accessions in (**a-c**); accessions were ordered according to the order in (**a**).

**Fig. S11 Feature interactions revealed by interpreting SHAP values of a feature in accessions with different values for another feature.** (**a-e**) T-T interactions of *SUPPRESSOR OF OVEREXPRESSION OF CO 1* (*SOC1*) with *FLOWERING LOCUS T* (*FT*) (**a**), MIR172B (**b**), *SQUAMOSA PROMOTER BINDING PROTEIN-LIKE 5* (*SPL5*) (**c**), *FLOWERING LOCUS C* (*FLC*) (**d**), and *PHYTOCHROME INTERACTING FACTOR 3* (*PIF3*) (**e**). (**f-g**) T-gbM interactions of SOC1 with *LOV KELCH PROTEIN 2* (*LKP2*) (**f**) and *FLAVIN-BINDING, KELCH REPEAT, F BOX 1* (*FKF1*) (**g**). (**h**) G-G interaction between *AT1G11520* and *HASTY 1* (*HST1*). (**i**) gbM-gbM interaction between *INDUCER OF CBF EXPRESSION 1* (*ICE1*) and VIN3-LIKE 3 (*VIL3*). (**j**) G-gbM interaction between *AT1G11520* and *FY*. The percentile rank of each feature interaction is listed above the corresponding figure. Alt: alternative allele; ref: reference allele. Equations are for regression of a given feature values (x-axis) and its SHAP values (y-axis) in accessions with higher (red) or lower (blue) values of the other feature (color legend, the cutoff is indicated by red and blue arrows).

**Fig. S12 Percentile thresholds used for inclusion of single site methylation data in model building.** (**a**) Distribution of methylation sites based on proportion of missing data across accessions. x-axis: number of accessions containing information for a given site; y-axis: frequency of sites that had a certain proportion of accessions with single-site methylation data available. Red, orange, and purple dashed lines: 50^th^, 75^th^, and 90^th^ percentiles of the distribution, respectively. (**b-c**) Accuracy of rrBLUP (**b**) and RF (**c**) flowering time prediction models on the test set using datasets with different thresholds. P/A: presence/absence of methylation at a cytosine site; Proportion: the proportion of methylated reads over all the reads mapped to a cytosine site. Numbers in the parentheses in (**c**) indicate the numbers of single site methylation values remaining in the analysis. (**d**) Diagram showing the bins of gene-based regions for summarizing methylation levels of single sites. A gene body region, defined as the region between the TSS (transcription state site) and TTS (transcription termination site) sites, is split into 20 bins; the upstream (500 bp upstream of the TSS) and downstream (500 bp downstream of the TTS) regions are each split into 5 bins.

**Fig. S13 Odds ratio between the number of identified benchmark flowering time genes and number of genes expected by random chance.** Odds ratio was from Fisher’s exact test, and values above 1 indicate that there were more benchmark genes identified than randomly expected when the 95^th^ (**a**) and 99^th^ (**b**) percentiles were used as the importance value thresholds. Coef: coefficient; imp: gini importance. SD: short day; LD: long day.

## Code availability

All the scripts and datasets used in this study are available on Github at: https://github.com/ShiuLab/Manuscript_Code/tree/master/2023_Ath_multi-omics

## Acknowledgements

We thank Elyse Vischulis for help in data collection, and Chad Niederhuth and Sunil Kumar Kenchanmane Raju for helpful discussions on DNA methylation. This work was supported by the U.S. Department of Energy Great Lakes Bioenergy Research Center (BER DE-SC0018409 to SHS), the National Science Foundation (DGE-1828149 and IOS-2218206 to SHS, KSA, and SL; IOS-2107215 and MCB-2210431 to MDL and SHS), and the Scientific Research Foundation of and the Major Scientific Research Tasks from Kunpeng Institute of Modern Agriculture at Foshan (KIMAQD2022003 and KIMA-ZDKY2022004 to PW).

## Author Contributions

PW and SHS conceived and designed the study with input from SL, KSA, and MDL. PW and SL performed the methylation-related analysis. PW and KSA analyzed the feature importance. PJK created the gene mutants, and MDL conducted the assessment of flowering time for loss-of-function mutant plants. PW conducted all other analyses. PW, SL, KSA, MDL, and SHS wrote the paper. All authors read and approved the final manuscript.

