## Supplementary material for "Prediction of plant complex traits via integration of multi-omics data": Fig. S1-13

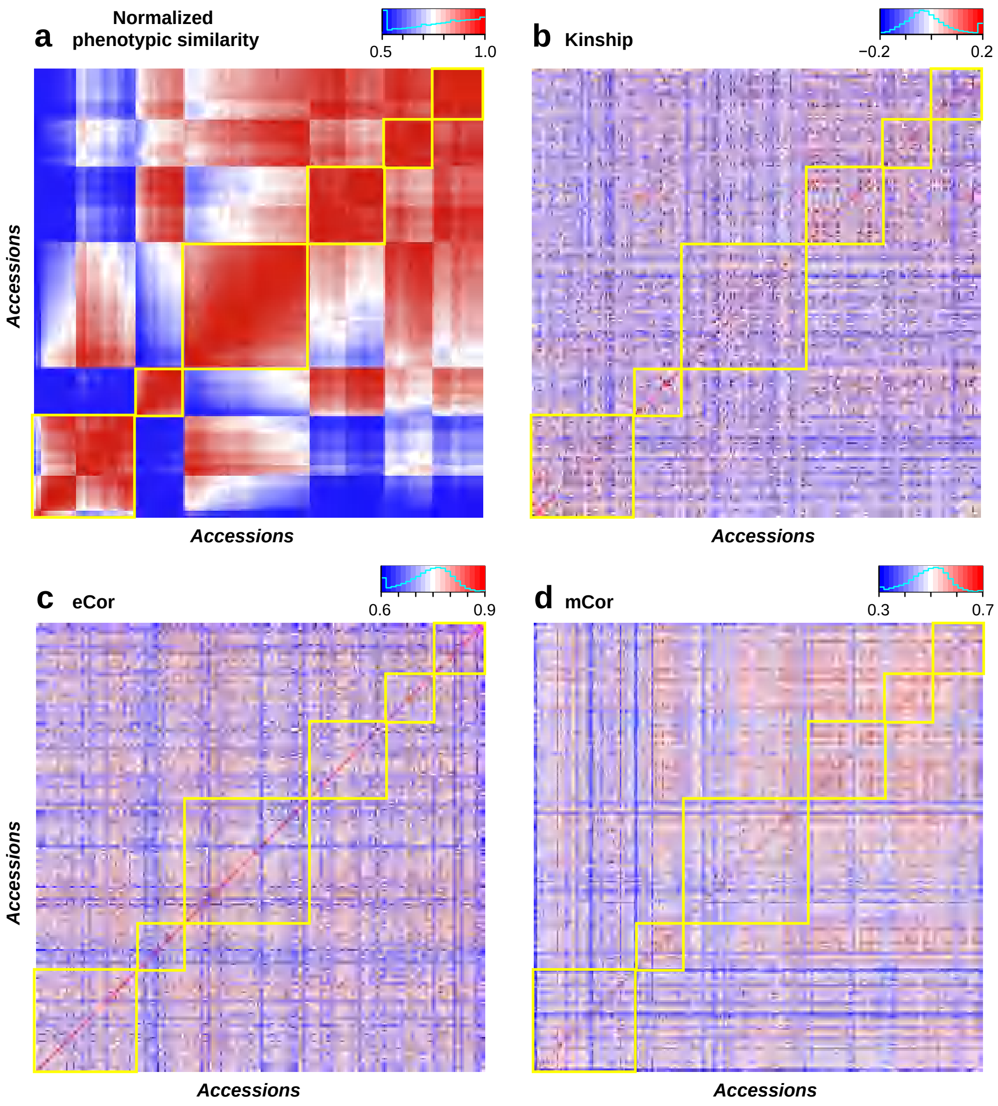

a
Normalized phenotypic similarity
0.5
1.0
b
Kinship
−0.2
0.2
Accessions
Accessions
Accessions
c
eCor
0.6
0.9
d
mCor
0.3
0.7
Accessions
Accessions
Accessions

**Fig. S2**

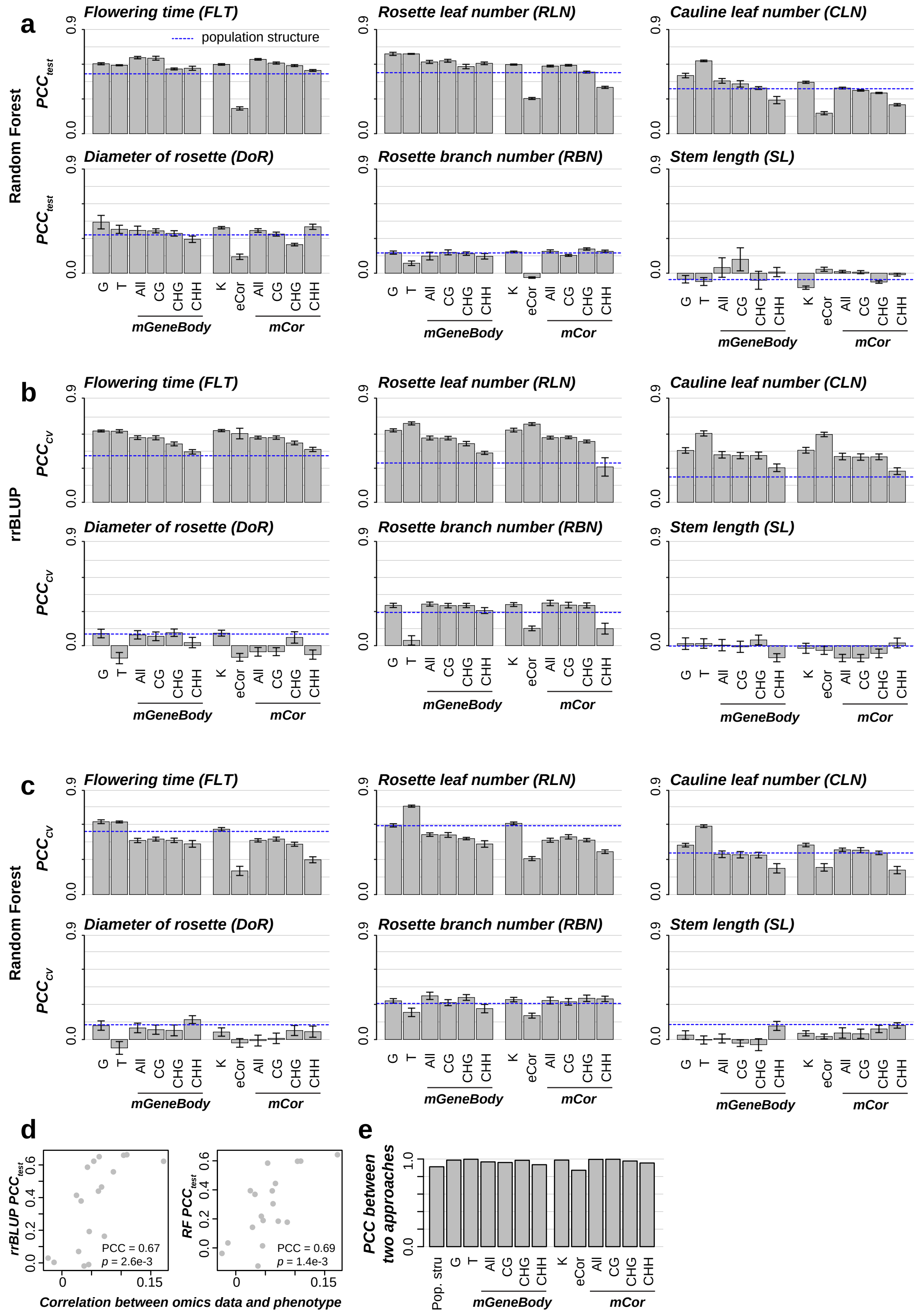

a
Random Forest
Flowering time (FLT)
population structure
Rosette leaf number (RLN)
Cauline leaf number (CLN)
Diameter of rosette (DoR)
Rosette branch number (RBN)
Stem length (SL)
b
rrBLUP
c
Random Forest
G
T
All
CG
CHG
CHH
K
eCor
mGeneBody
mCor
0.0
0.9
d
PCC = 0.67
p = 2.6e-3
PCC = 0.69
p = 1.4e-3
Correlation between omics data and phenotype
e
PCC between two approaches
Pop. stru

Fig. S3

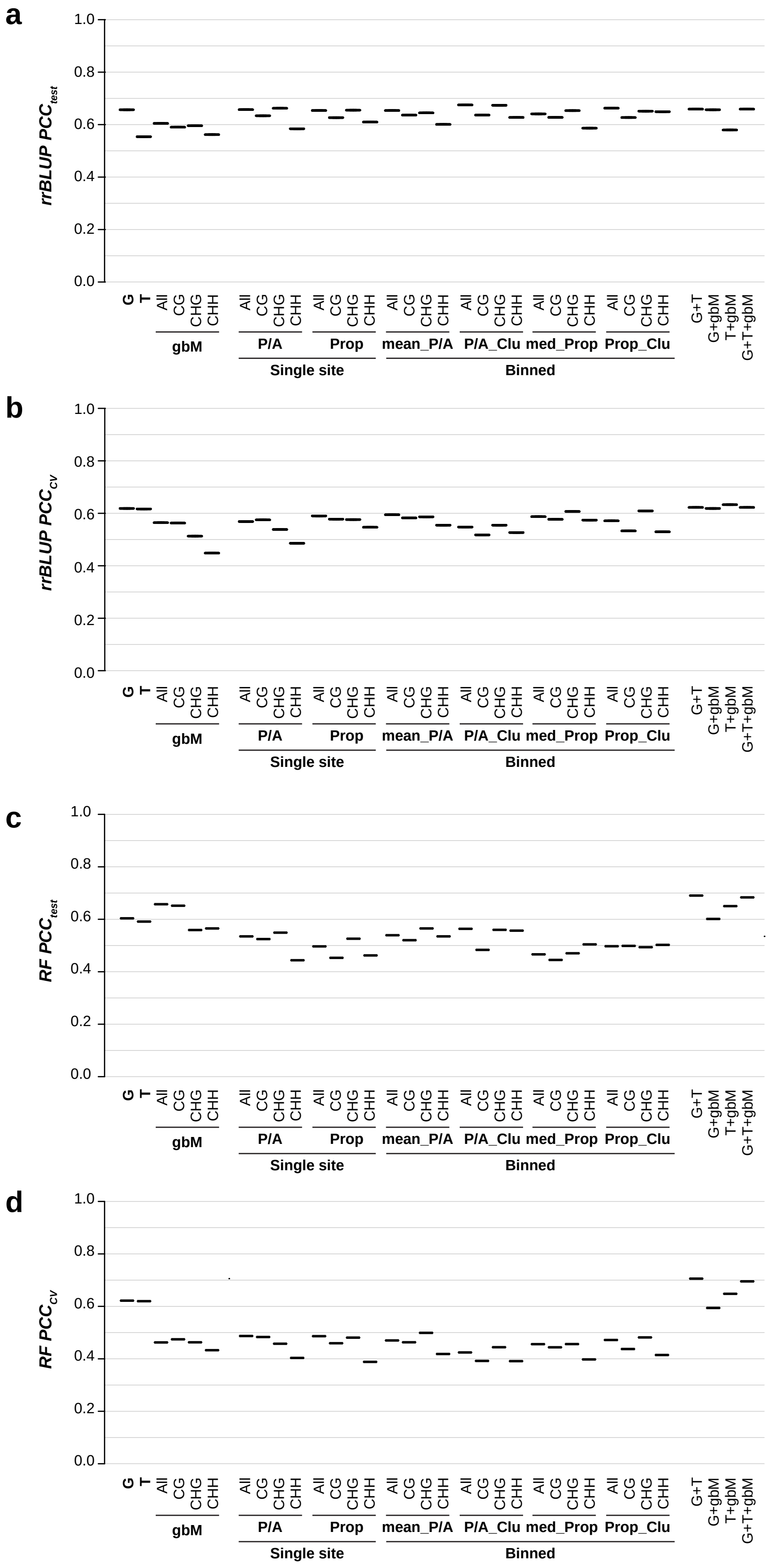
a
b
c
d
rrBLUP PCC test
rrBLUP PCC cv
RF PCC test
RF PCC cv
1.0
0.8
0.6
0.4
0.2
0.0
G
T
All
CG
CHG
CHH
gbM
P/A
Prop
mean_P/A
P/A_Clu
med_Prop
Prop_Clu
Single site
Binned
G+T
G+gbM
T+gbM
G+T+gbM

Fig. S4

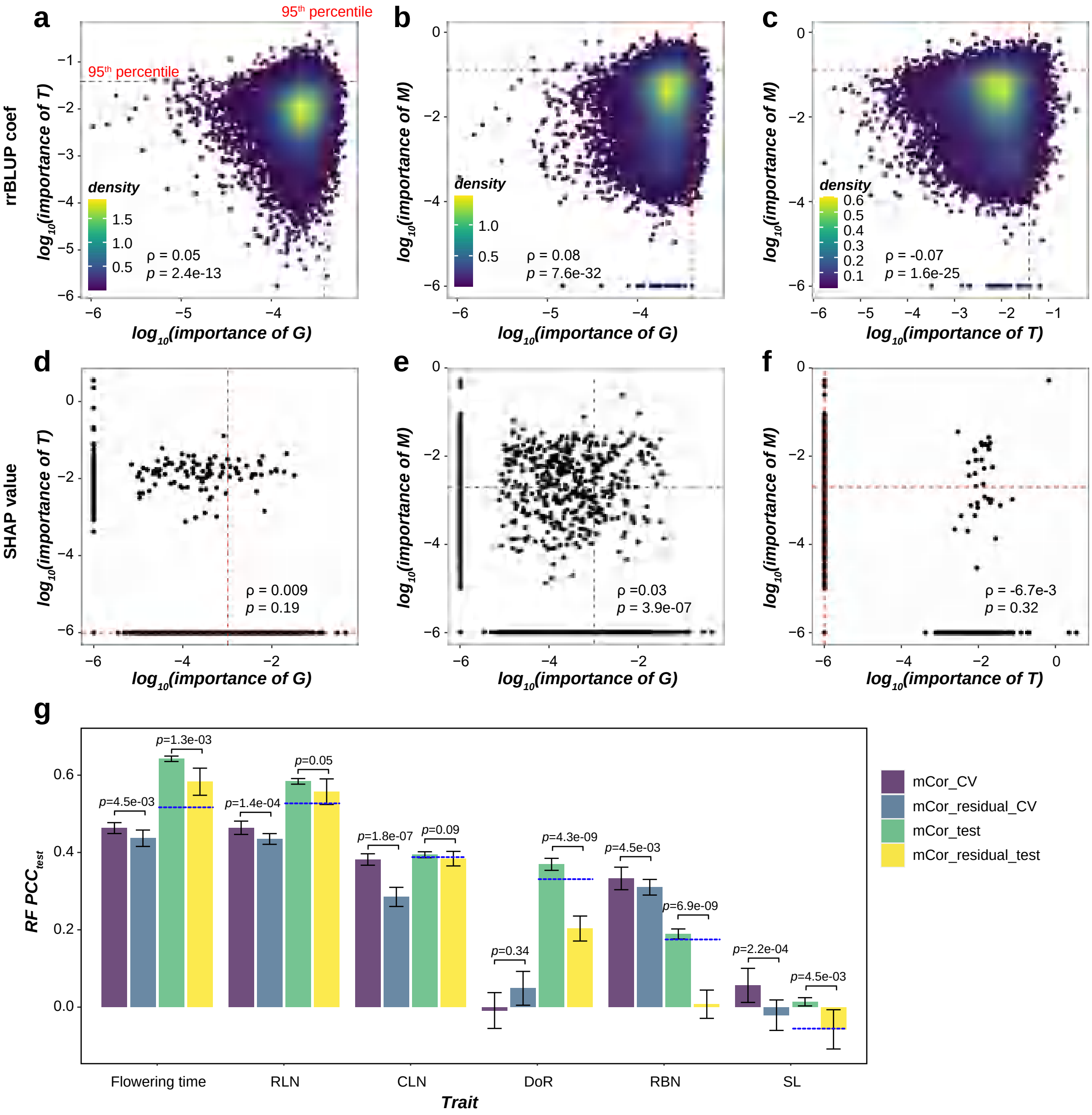
a
b
c
d
e
f
g
95th percentile
rrBLUP coef
log10(importance of T)
log10(importance of G)
log10(importance of M)
log10(importance of T)
density
ρ = 0.05
p = 2.4e-13
ρ = 0.08
p = 7.6e-32
ρ = -0.07
p = 1.6e-25
SHAP value
ρ = 0.009
p = 0.19
ρ =0.03
p = 3.9e-07
ρ = -6.7e-3
p = 0.32
RF PCCtest
p=4.5e-03
p=1.3e-03
p=1.4e-04
p=0.05
p=1.8e-07
p=0.09
p=0.34
p=4.3e-09
p=4.5e-03
p=6.9e-09
p=2.2e-04
p=4.5e-03
Flowering time
RLN
CLN
DoR
RBN
SL
Trait
mCor_CV
mCor_residual_CV
mCor_test
mCor_residual_test

**Fig. S5**

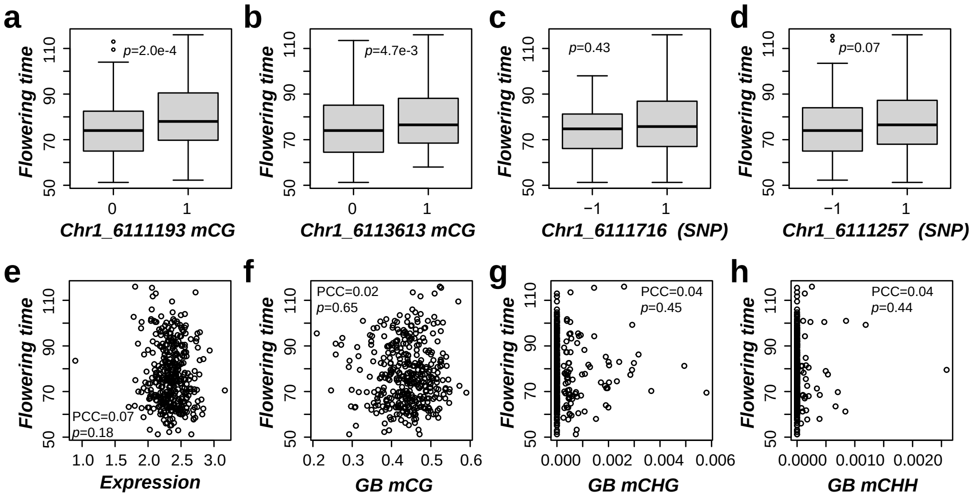
a
b
c
d
p=2.0e-4
p=4.7e-3
p=0.43
p=0.07
Flowering time
Chr1_6111193 mCG
Chr1_6113613 mCG
Chr1_6111716 (SNP)
Chr1_6111257 (SNP)
e
f
g
h
PCC=0.07
p=0.18
PCC=0.02
p=0.65
PCC=0.04
p=0.45
PCC=0.04
p=0.44
Expression
GB mCG
GB mCHG
GB mCHH

#### Fig. S6

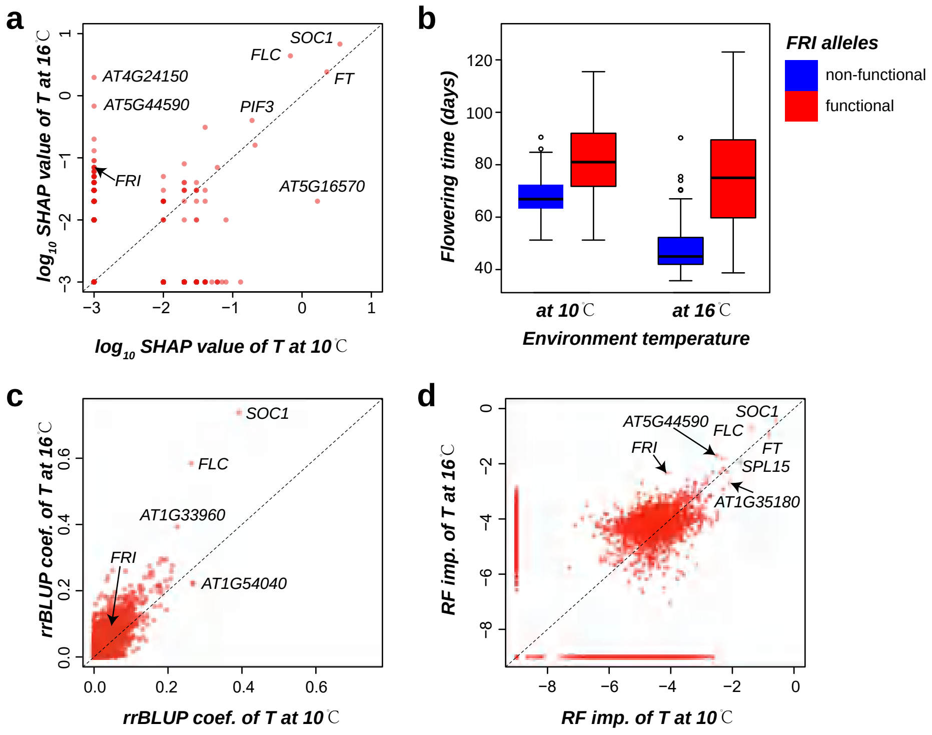
a
SOC1
FLC
FT
AT4G24150
AT5G44590
PIF3
FRI
AT5G16570
log10 SHAP value of T at 16℃
log10 SHAP value of T at 10℃
b
FRI alleles
non-functional
functional
Flowering time (days)
at 10℃
at 16℃
Environment temperature
c
SOC1
FLC
AT1G33960
FRI
AT1G54040
rrBLUP coef. of T at 16℃
rrBLUP coef. of T at 10℃
d
AT5G44590
SOC1
FLC
FRI
FT
SPL15
AT1G35180
RF imp. of T at 16℃
RF imp. of T at 10℃

#### Fig. S7

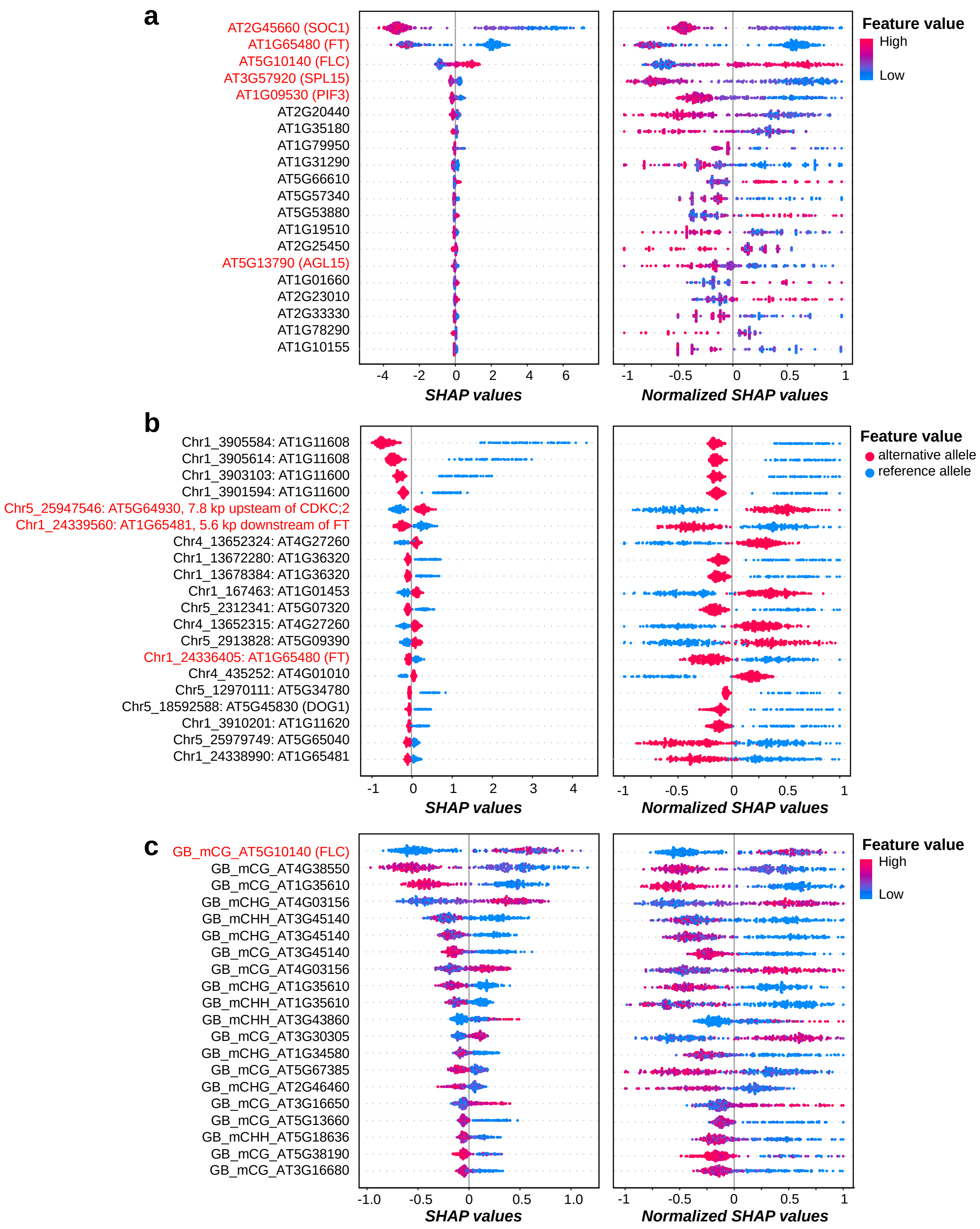
a
AT2G45660 (SOC1)
AT1G65480 (FT)
AT5G10140 (FLC)
AT3G57920 (SPL15)
AT1G09530 (PIF3)
AT2G20440
AT1G35180
AT1G79950
AT1G31290
AT5G66610
AT5G57340
AT5G53880
AT1G19510
AT2G25450
AT5G13790 (AGL15)
AT1G01660
AT2G23010
AT2G33330
AT1G78290
AT1G10155
SHAP values
Normalized SHAP values
Feature value
High
Low
b
Chr1_3905584: AT1G11608
Chr1_3905614: AT1G11608
Chr1_3903103: AT1G11600
Chr1_3901594: AT1G11600
Chr5_25947546: AT5G64930, 7.8 kp upsteam of CDKC;2
Chr1_24339560: AT1G65481, 5.6 kp downstream of FT
Chr4_13652324: AT4G27260
Chr1_13672280: AT1G36320
Chr1_13678384: AT1G36320
Chr1_167463: AT1G01453
Chr5_2312341: AT5G07320
Chr4_13652315: AT4G27260
Chr5_2913828: AT5G09390
Chr1_24336405: AT1G65480 (FT)
Chr4_435252: AT4G01010
Chr5_12970111: AT5G34780
Chr5_18592588: AT5G45830 (DOG1)
Chr1_3910201: AT1G11620
Chr5_25979749: AT5G65040
Chr1_24338990: AT1G65481
SHAP values
Normalized SHAP values
Feature value
alternative allele
reference allele
c
GB_mCG_AT5G10140 (FLC)
GB_mCG_AT4G38550
GB_mCG_AT1G35610
GB_mCHG_AT4G03156
GB_mCHH_AT3G45140
GB_mCHG_AT3G45140
GB_mCG_AT3G45140
GB_mCG_AT4G03156
GB_mCHG_AT1G35610
GB_mCHH_AT1G35610
GB_mCHH_AT3G43860
GB_mCG_AT3G30305
GB_mCHG_AT1G34580
GB_mCG_AT5G67385
GB_mCHG_AT2G46460
GB_mCG_AT3G16650
GB_mCG_AT5G13660
GB_mCHH_AT5G18636
GB_mCG_AT5G38190
GB_mCG_AT3G16680
SHAP values
Normalized SHAP values
Feature value
High
Low

#### Fig. S8

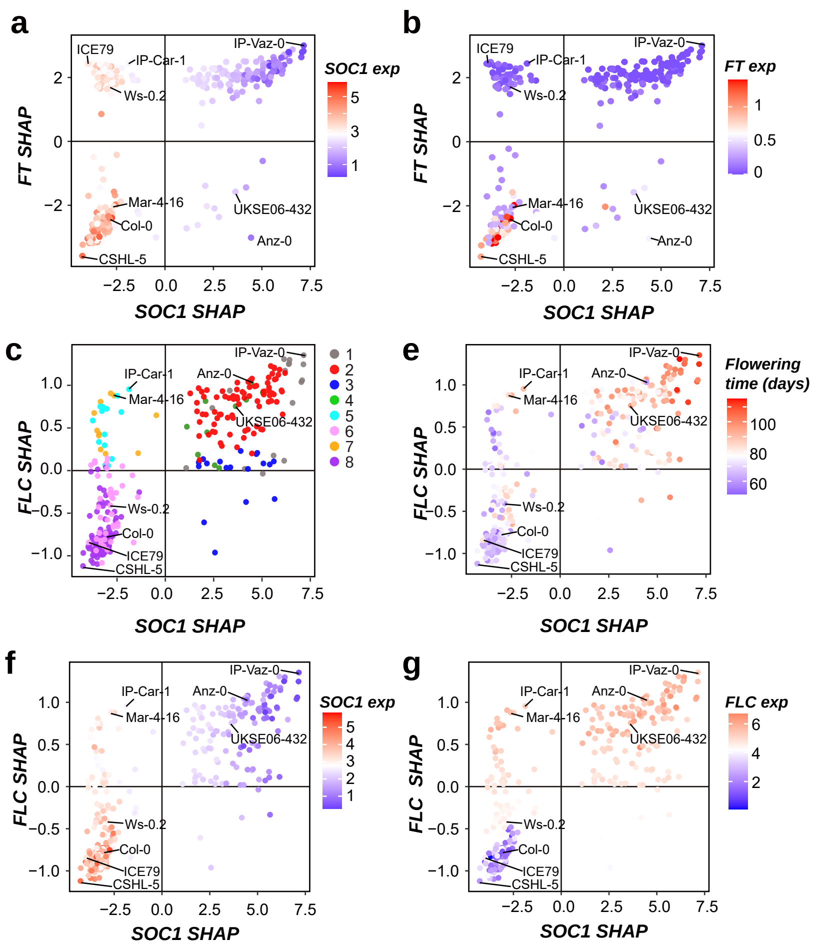
a
b
c
e
f
g
SOC1 SHAP
FT SHAP
FLC SHAP
SOC1 exp
FT exp
FLC exp
Flowering time (days)
ICE79
IP-Car-1
Ws-0.2
IP-Vaz-0
Mar-4-16
Col-0
CSHL-5
UKSE06-432
Anz-0

#### Fig. S9

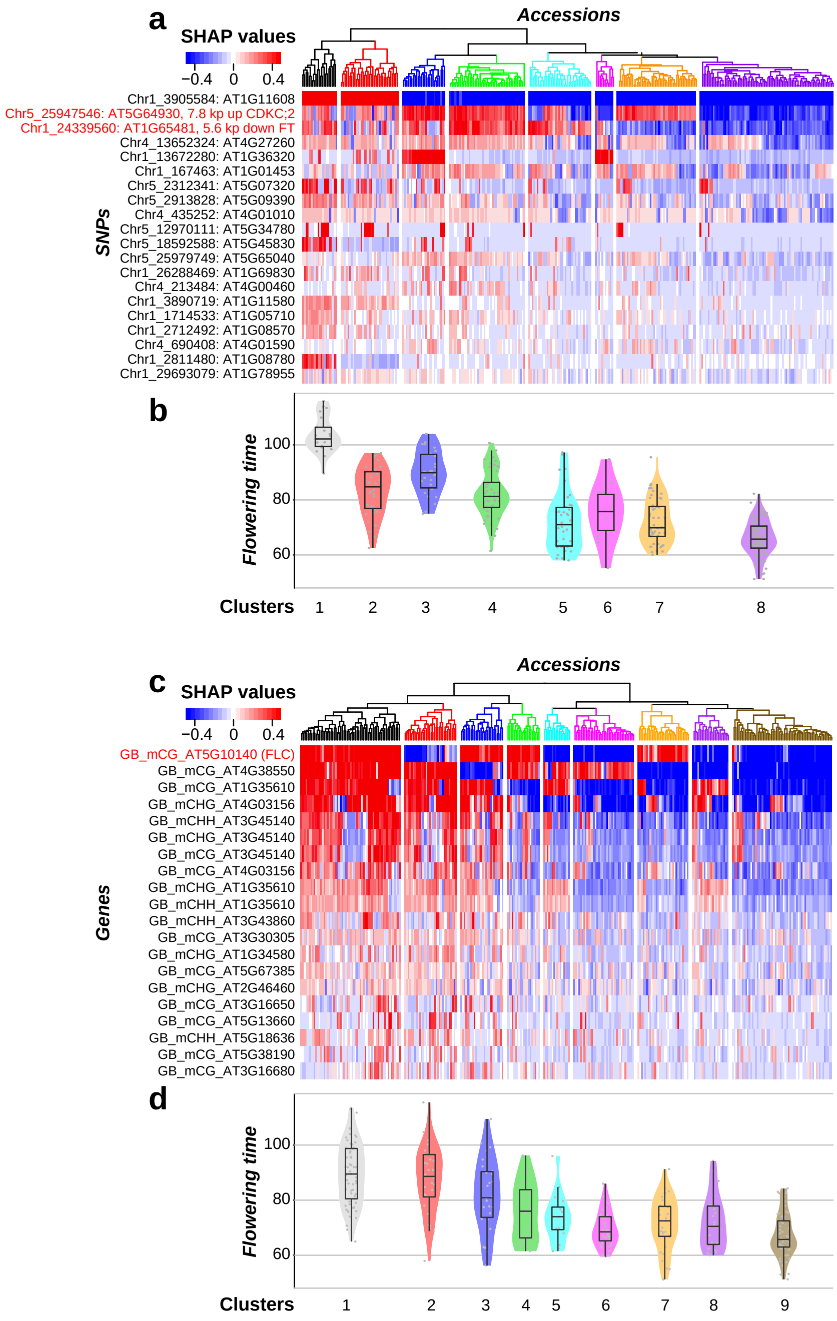
a
SHAP values
−0.4 0 0.4
Accessions
SNPs
Chr1_3905584: AT1G11608
Chr5_25947546: AT5G64930, 7.8 kp up CDKC;2
Chr1_24339560: AT1G65481, 5.6 kp down FT
Chr4_13652324: AT4G27260
Chr1_13672280: AT1G36320
Chr1_167463: AT1G01453
Chr5_2312341: AT5G07320
Chr5_2913828: AT5G09390
Chr4_435252: AT4G01010
Chr5_12970111: AT5G34780
Chr5_18592588: AT5G45830
Chr5_25979749: AT5G65040
Chr1_26288469: AT1G69830
Chr4_213484: AT4G00460
Chr1_3890719: AT1G11580
Chr1_1714533: AT1G05710
Chr1_2712492: AT1G08570
Chr4_690408: AT4G01590
Chr1_2811480: AT1G08780
Chr1_29693079: AT1G78955
b
Flowering time
100
80
60
Clusters
1 2 3 4 5 6 7 8
c
SHAP values
−0.4 0 0.4
Accessions
Genes
GB_mCG_AT5G10140 (FLC)
GB_mCG_AT4G38550
GB_mCG_AT1G35610
GB_mCHG_AT4G03156
GB_mCHH_AT3G45140
GB_mCHG_AT3G45140
GB_mCG_AT3G45140
GB_mCG_AT4G03156
GB_mCHG_AT1G35610
GB_mCHH_AT1G35610
GB_mCHH_AT3G43860
GB_mCG_AT3G30305
GB_mCHG_AT1G34580
GB_mCG_AT5G67385
GB_mCHG_AT2G46460
GB_mCG_AT3G16650
GB_mCG_AT5G13660
GB_mCHH_AT5G18636
GB_mCG_AT5G38190
GB_mCG_AT3G16680
d
Flowering time
100
80
60
Clusters
1 2 3 4 5 6 7 8 9

#### Fig. S10

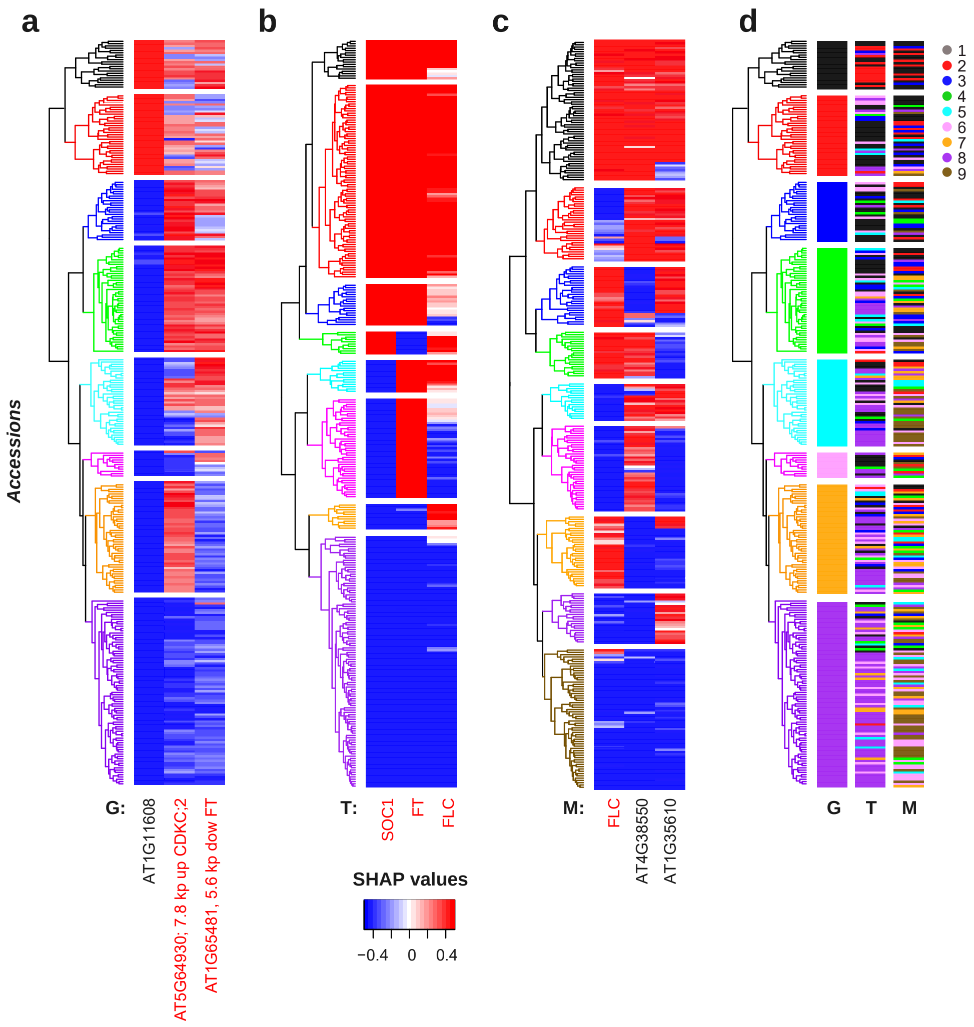
a
b
c
d
Accessions
G:
AT1G11608
AT5G64930; 7.8 kp up CDKC;2
AT1G65481, 5.6 kp dow FT
T:
SOC1
FT
FLC
M:
FLC
AT4G38550
AT1G35610
G
T
M
1
2
3
4
5
6
7
8
9
SHAP values
−0.4
0
0.4

### Fig. S11

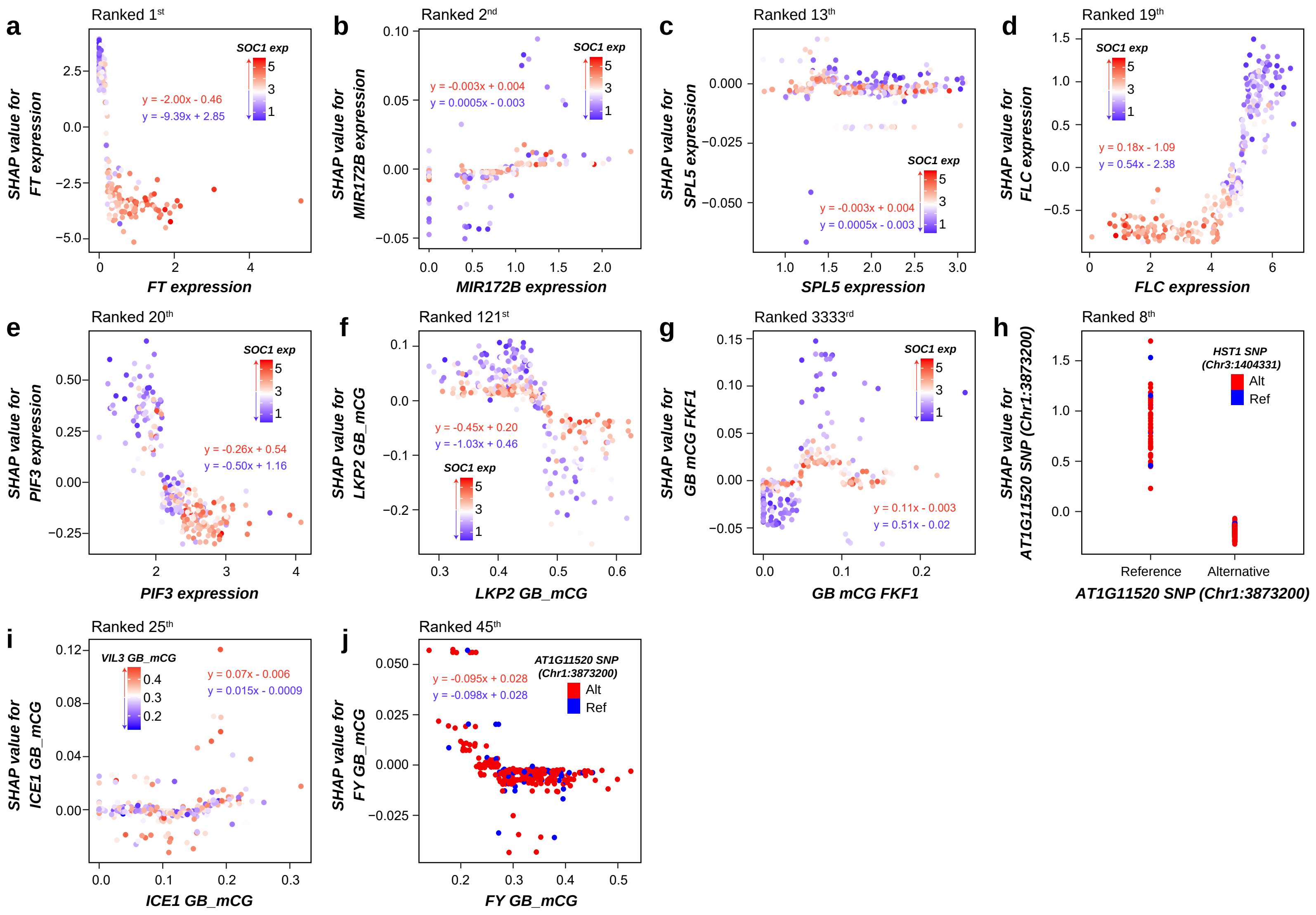

a
Ranked 1st
SOC1 exp
y = -2.00x - 0.46
y = -9.39x + 2.85
SHAP value for FT expression
FT expression
b
Ranked 2nd
SOC1 exp
y = -0.003x + 0.004
y = 0.0005x - 0.003
SHAP value for MIR172B expression
MIR172B expression
c
Ranked 13th
SOC1 exp
y = -0.003x + 0.004
y = 0.0005x - 0.003
SHAP value for SPL5 expression
SPL5 expression
d
Ranked 19th
SOC1 exp
y = 0.18x - 1.09
y = 0.54x - 2.38
SHAP value for FLC expression
FLC expression
e
Ranked 20th
SOC1 exp
y = -0.26x + 0.54
y = -0.50x + 1.16
SHAP value for PIF3 expression
PIF3 expression
f
Ranked 121st
y = -0.45x + 0.20
y = -1.03x + 0.46
SOC1 exp
SHAP value for LKP2 GB_mCG
LKP2 GB_mCG
g
Ranked 3333rd
SOC1 exp
y = 0.11x - 0.003
y = 0.51x - 0.02
SHAP value for GB mCG FKF1
GB mCG FKF1
h
Ranked 8th
HST1 SNP (Chr3:1404331)
Alt
Ref
SHAP value for AT1G11520 SNP (Chr1:3873200)
Reference
Alternative
AT1G11520 SNP (Chr1:3873200)
i
Ranked 25th
VIL3 GB_mCG
y = 0.07x - 0.006
y = 0.015x - 0.0009
SHAP value for ICE1 GB_mCG
ICE1 GB_mCG
j
Ranked 45th
y = -0.095x + 0.028
y = -0.098x + 0.028
AT1G11520 SNP (Chr1:3873200)
Alt
Ref
SHAP value for FY GB_mCG
FY GB_mCG

#### Fig. S12

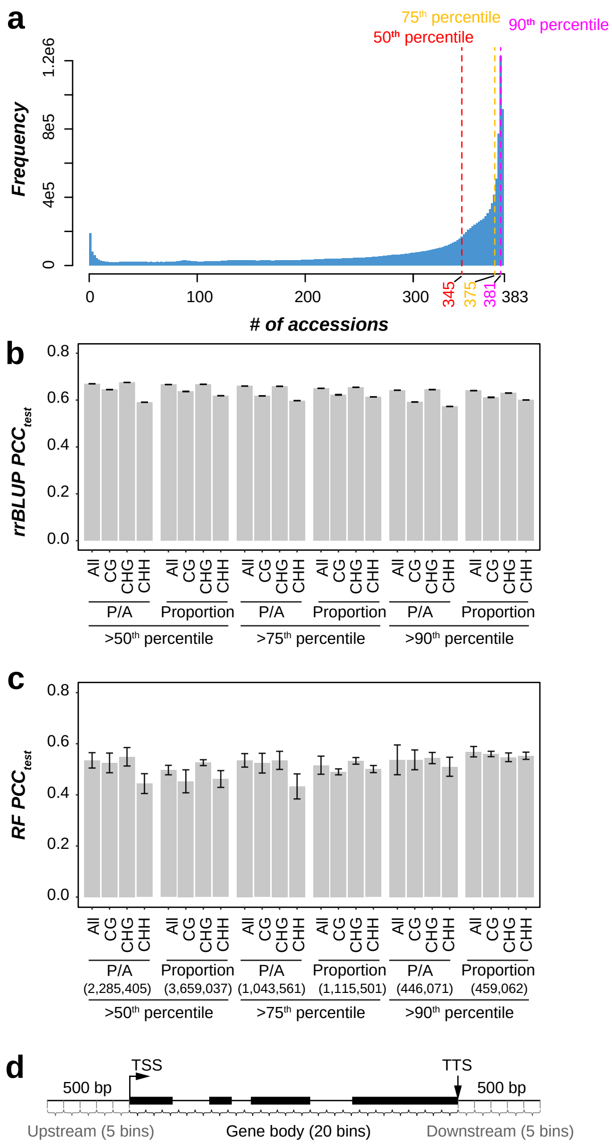
a
75th percentile
50th percentile
90th percentile
Frequency
0
4e5
8e5
1.2e6
0
100
200
300
345
375
381
383
### of accessions
b
rrBLUP PCC_test
0.0
0.2
0.4
0.6
0.8
All CG CHG CHH
P/A
Proportion
>50th percentile
>75th percentile
>90th percentile
c
RF PCC_test
P/A (2,285,405)
Proportion (3,659,037)
P/A (1,043,561)
Proportion (1,115,501)
P/A (446,071)
Proportion (459,062)
d
TSS
TTS
500 bp
500 bp
Upstream (5 bins)
Gene body (20 bins)
Downstream (5 bins)

### Fig. S13

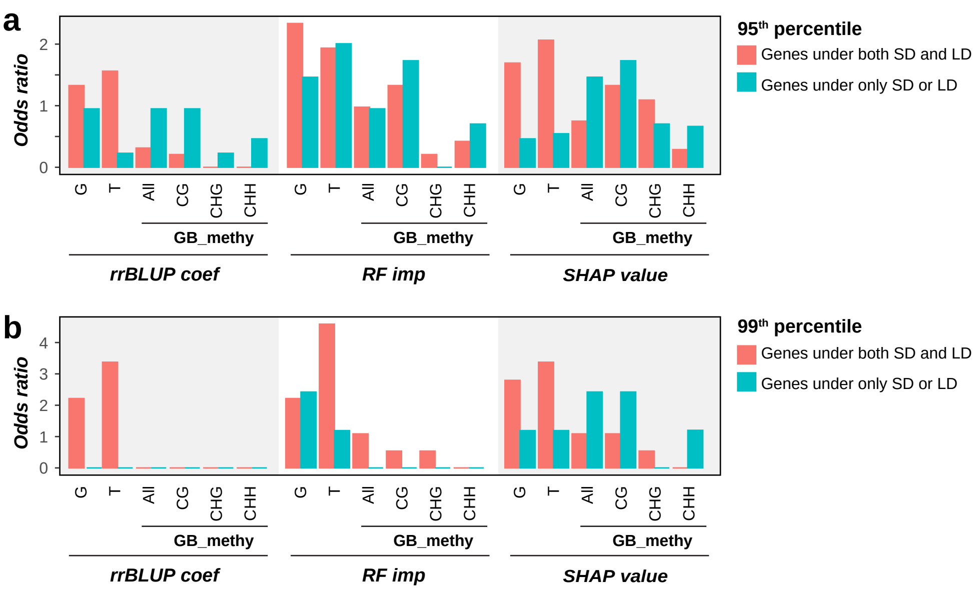
a
Odds ratio
0
1
2
G
T
All
CG
CHG
CHH
GB_methy
rrBLUP coef
RF imp
SHAP value
95th percentile
Genes under both SD and LD
Genes under only SD or LD
b
Odds ratio
0
1
2
3
4
G
T
All
CG
CHG
CHH
GB_methy
rrBLUP coef
RF imp
SHAP value
99th percentile
Genes under both SD and LD
Genes under only SD or LD
